# Noncanonical scaffolding of G_αi_ and β-arrestin by G protein-coupled receptors

**DOI:** 10.1101/629576

**Authors:** Jeffrey S. Smith, Thomas F. Pack, Asuka Inoue, Claudia Lee, Xinyu Xiong, Kevin Zheng, Alem W. Kahsai, Issac Choi, Zhiyuan Ma, Ian M. Levitan, Lauren K. Rochelle, Dean P. Staus, Joshua C. Snyder, Marc G. Caron, Sudarshan Rajagopal

**Author notes:** These authors contributed equally to this work (Jeffrey S. Smith, Thomas F. Pack).

## Abstract

G-protein-coupled receptors (GPCRs) enable cells to sense and respond appropriately to hormonal and environmental signals, and are a target of ~30% of all FDA-approved medications. Canonically, each GPCR couples to distinct G_α_ proteins, such as G_αs_, G_αi_, G_αq_ or G_α12/13_, as well as β-arrestins. These transducer proteins translate and integrate extracellular stimuli sensed by GPCRs into intracellular signals through what are broadly considered separable signalling pathways. However, the ability of G_α_ proteins to directly interact with β-arrestins to integrate signalling has not previously been appreciated. Here we show a novel interaction between G_αi_ protein family members and β-arrestin. G_αi_:β-arrestin complexes were formed by all GPCRs tested, regardless of their canonical G protein isoform coupling, and could bind both GPCRs as well as the extracellular signal-regulated kinase (ERK). This novel paradigm of G_αi_:β-arrestin scaffolds enhances our understanding of GPCR signalling.

## Introduction

Classically, G protein-coupled receptors (GPCRs) couple to a distinct G_α_ protein and activate G proteins by catalyzing guanine nucleotide exchange. β-arrestins were subsequently discovered to both regulate and transduce GPCR signalling. More recently, the ability of G proteins^1^ and β-arrestins^2^ to coordinate their signalling has been suggested by the demonstration of “megaplex” signalling complexes consisting of a GPCR together with *both* G protein and β-arrestin^3,4^. However, the ability of G_α_ proteins to *directly* interact with β-arrestins to form signalling scaffolds has not previously been appreciated. Here, we use a ‘complex’ bioluminescent resonance energy transfer (BRET) approach to identify a novel G_α_ protein-β-arrestin interaction. Agonist treatment of the G_αs_-coupled vasopressin type 2 receptor (V_2_R) paradoxically catalysed the formation of G_αi_:β-arrestin scaffolds that were not observed with other G_α_ families, despite the inability of the vasopressin-treated receptor to promote canonical G_αi_-mediated signalling. These G_αi_:β-arrestin complexes were also formed downstream of all GPCRs tested, including the β_2_-adrenergic receptor (β_2_AR), neurotensin receptor type 1 (NT_1_R), dopamine D1 and D2 receptors (D_1_R, D_2_R), and C-X-C motif chemokine receptor 3 (CXCR3), of which only D_2_R and CXCR3 canonically activate G_αi_. These scaffolds were not observed to form with other G_α_ subtypes. The G_αi_:β-arrestin scaffolds can form “megaplexes” with GPCRs and can also bind extracellular signal-regulated kinase (ERK). Disrupting G_αi_ and β-arrestin interactions eliminated V_2_R mediated transduction of ERK phosphorylation. In addition, disrupting G_αi_ and β-arrestin interactions eliminated GPCR-mediated migration in response to a β-arrestin-biased agonist that does not stimulate canonical G_αi_ signalling. These results uncover a novel GPCR signalling paradigm involving the formation of noncanonical G_αi_:β-arrestin signalling scaffolds.

### β-arrestin, G_αi_, and receptor form complexes

It is well established that GPCRs differentially associate with β-arrestins following agonist treatment^5^. For example, agonist treatment of certain receptors, such as the G_αs_-coupled V_2_R, results in a long-lived receptor association with β-arrestin, contrasting with other receptors, such as the β_2_AR, that form transient interactions with β-arrestin with dissociation occurring at or near the plasma membrane^6^. More recently, it has been shown that G_αs_, β-arrestin, and a GPCR can form signalling ‘megaplexes’^3^. To further interrogate the composition of megaplexes, we utilized a ‘complex BRET’ approach (Fig. 1a), similar to other BRET-based strategies to assess complex formation^7,8^, to confirm simultaneous interactions between G_αs_, β-arrestin, and V_2_R following agonist treatment (Fig. 1b,c). Complex BRET requires complementation of a low affinity split luciferase (nanoBiT) (shown not to affect underlying protein:protein interactions^9^) by complementing a small peptide (smBiT) fused to one protein to a large protein fragment (LgBiT) fused to another protein of interest. The signal generated by complementation of this split luciferase can then transfer to a third protein tagged with a fluorescent protein acceptor, monomeric Kusabira Orange (mKO), generating a BRET response. Thus, this technique enables real-time quantification of interactions between a two-protein complex and a third protein in living cells.

**Figure 1:**
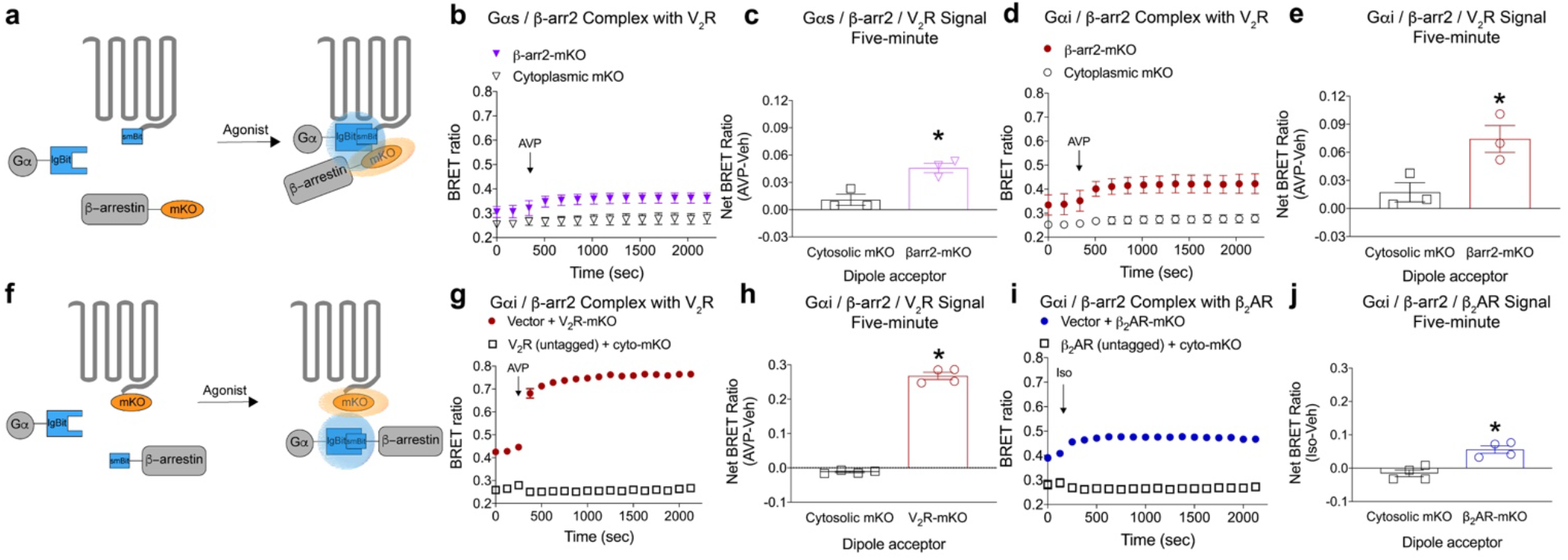
Formation of G protein:β-arrestin:GPCR megaplexes. **a**, Arrangement of luciferase fragments and mKO acceptor fluorophore for complex BRET on G protein (LgBiT), β-arrestin (mKO), and V_2_R (smBiT). HEK 293T cells were transiently transfected with the indicated receptor and assay components and stimulated with the indicated agonist or vehicle. **b**, Complex BRET ratio of G_αs_:β-arrestin:V_2_R following AVP (500 nM) treatment. After AVP treatment, an increase in the BRET ratio was observed in cells expressing β-arrestin-mKO, but not cytosolic mKO. **c**, Quantification of G_αs_:mKO:V_2_R complex formation in cells treated with either vehicle or AVP at a single five-minute timepoint. Full kinetic data is available in the extended data. **d**, Similar experiment to panel b, except testing the ability of G_αi_ to form a ‘megaplex.’ Complex BRET ratio of G_αi_:β-arrestin:V_2_R following treatment with AVP. After AVP treatment, an increase in the BRET ratio was observed in cells expressing β-arrestin-mKO, but not cytosolic mKO, which is similar to panel **b. e**, Quantification of the G_αi_:mKO:V_2_R complex formation in cells treated with either vehicle or AVP at a single five-minute timepoint. **f**, Rearrangement of complex BRET components on G protein (LgBiT), β-arrestin (SmBiT), and V_2_R (mKO). **g**, Complex BRET ratio of G_αi_:β-arrestin:V_2_R following AVP treatment. Rearrangement of complex BRET tags increased the observed signal when compared to panel **d. h**, Five-minute quantification of G_αi_-arrestin:V_2_R complexes relative to vehicle treatment. **i**, Similar experiment to panel **g**, except testing the ability of G_αi_:β-arrestin to form a megaplex with the β_2_AR as opposed to the V_2_R. After isoproterenol (10 μM) treatment, an increase in the BRET ratio was observed in cells expressing β_2_AR-mKO, but not cytosolic mKO. **j**, Five-minute quantification of G_αi_:β-arrestin:mKO complexes induced by isoproterenol relative vehicle treatment. For kinetic experiments, \**P*<0.05 by two-way ANOVA, Fischer’s post hoc analysis with a significant difference between treatments; for five-minute quantification, \**P*<0.05 by student’s two-tailed t-test; for TSA \**P*<0.05 with Bonferroni post hoc analysis. Panels b-e, n=3 per condition; panels g-j, n=4 per condition. Graphs show mean ± s.e.m. Cyto, cytoplasmic.

Using this technology, we were surprised to discover that the canonically G_αs_-coupled V_2_R also formed a ‘megaplex’ with G_αi_ and β-arrestin following agonist treatment (Fig. 1d,e). To further interrogate the G_αi_:β-arrestin:V_2_R megaplex, we proceeded to swap the location of dipole donor and acceptor components (Fig. 1f). Altering the location of complex BRET components increased the observed signal of the G_αi_-containing megaplex following agonist treatment, and further confirmed that G_αi_:β-arrestin complexes can associate with the V_2_R (Fig 1g,h, Extended Data Fig. 1a). Agonist treatment of the canonically G_αs_-coupled β_2_AR also formed G_αi_:β-arrestin:β_2_AR megaplexes (Fig 1i,j, Extended Data Fig. 1b), although less robustly than the V_2_R. We further validated the specificity of megaplex formation by simultaneously transfecting both mKO-tagged and untagged V_2_R and β_2_AR and treating with agonist. Only treating a mKO-tagged receptor with its cognate agonist formed an observable G_αi_:β-arrestin:GPCR megaplex (Extended Data Fig 1c-h), indicating a specific interaction and not a bystander effect.

We next confirmed formation of G_αi_:β-arrestin:V_2_R megaplexes in reconstituted and overexpressed systems. We first used purified megaplex components *in vitro* in a thermal stability assay (TSA), which measures conformational stability of proteins upon thermal denaturation^10^. A change in the melting profile of a given protein in the presence of another molecule relative to its appropriate reference is an indication of binding/formation of a new complex. Consistent with the G_αi_:β-arrestin:V_2_R interaction observed with complex BRET, a change in the melting profile was observed when recombinant G_αi_-megaplex components were combined in the presence of a stabilizing antibody fragment, Fab30^11^ (Extended Data Fig. 2). In additional support of G_αi_:β-arrestin:V_2_R megaplexes, immunoprecipitation of β-arrestin yielded both V_2_R and G_αi_ association when overexpressed in HEK 293 cells (Extended Data Fig. 3).

### G_αi_:β-arrestin:V_2_R complexes form at the plasma membrane

We then visualized colocalization of G_αi_, β-arrestin, and V_2_R using confocal microscopy (Fig. 2a). We validated the imaging parameters using single-colour controls to ensure accurate quantification of each component channel (Extended Data Fig. 4). Colocalization of G_αi_, β-arrestin, and V_2_R occurred after agonist treatment and was most prominent at the plasma membrane. Line scan analyses demonstrated plasma membrane-localized puncta consisting of each megaplex component after 5 minutes of agonist treatment (Fig. 2b,c). Thirty minutes after agonist treatment, clear endosomal β-arrestin:V_2_R colocalization was observed that lacked substantial G_αi_ (Fig. 2d). These observations suggest that formation of G_αi_:β-arrestin:V_2_R megaplexes occurs after agonist treatment and is most prominent at the plasma membrane.

**Figure 2:**
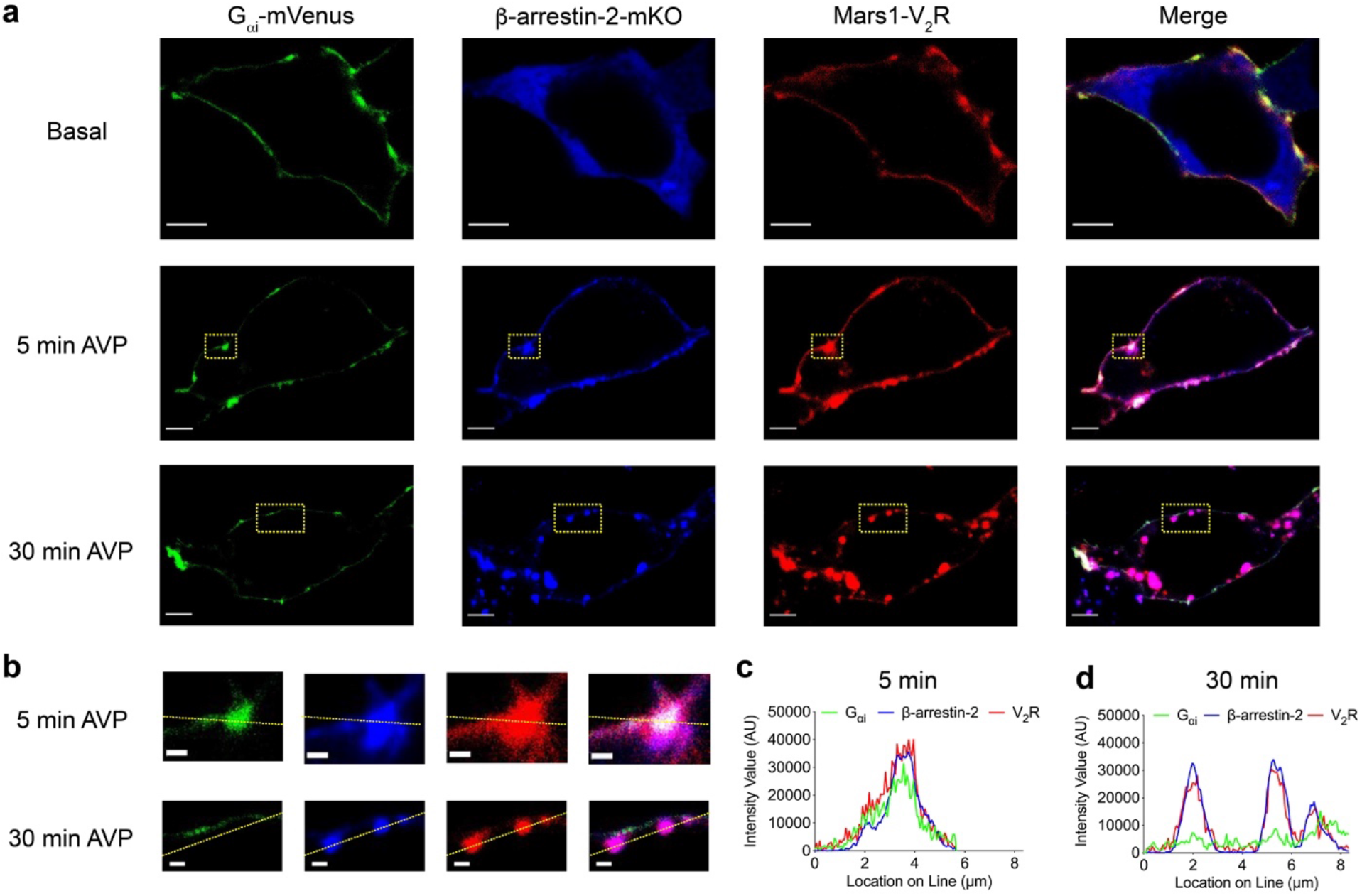
Confocal microscopy of G_αi_:β-arrestin:V_2_R complexes. Confocal microscopy analysis of AVP-induced complexes of G_αi_:β-arrestin:V_2_R in HEK 293 cells transfected with mVenus-tagged G_αi_, mKO-tagged β-arrestin-2 and Mars1-tagged V_2_R **a**, preceding treatment (basal), at 5 min, or at 30 min. Substantial co-localization of G_αi_:β-arrestin:V_2_R was observed at 5 min, with less appreciated at 30 min. **b**, inset of images in (**a**), scale bars, 1 μm. **c**, line scan analysis of 5-minute time point, demonstrating colocalization of fluorophores following AVP treatment. **d**, line-scan analysis of 30-minute time point. Scale bars, 5 μm. Data is representative of ten (basal), twenty (5 min) or fifteen (30 min) fields of view from three independent experiments. AVP was used at a concentration of 100 nM.

### G_αi_ forms a complex with β-arrestin following GPCR agonist treatment

The difference in magnitude of signal observed in complex BRET between G_αi_ and G_αs_ megaplexes suggested distinct interaction orientations in these complexes. It was recently shown that sustained G protein signalling exists at receptors following β-arrestin-dependent internalization^12^ and that β-arrestins are catalytically activated by an agonist-occupied receptor^13^. We therefore hypothesized that other critical interactions between G proteins and β-arrestins could be catalysed by agonist treatment of the V_2_R. We confirmed that the V_2_R primarily signals via G_αs_ (Fig. 3a) and recruits β-arrestin (Fig. 3b) following agonist treatment. Notably, V_2_R did not canonically signal via G_αi_, even under these overexpressed conditions (Fig. 3a). Similarly, we observed predominant G_αs_ signalling and β-arrestin recruitment at the β_2_AR, another canonically G_αs_-coupled receptor (Extended data Fig 5a,b).

**Figure 3:**
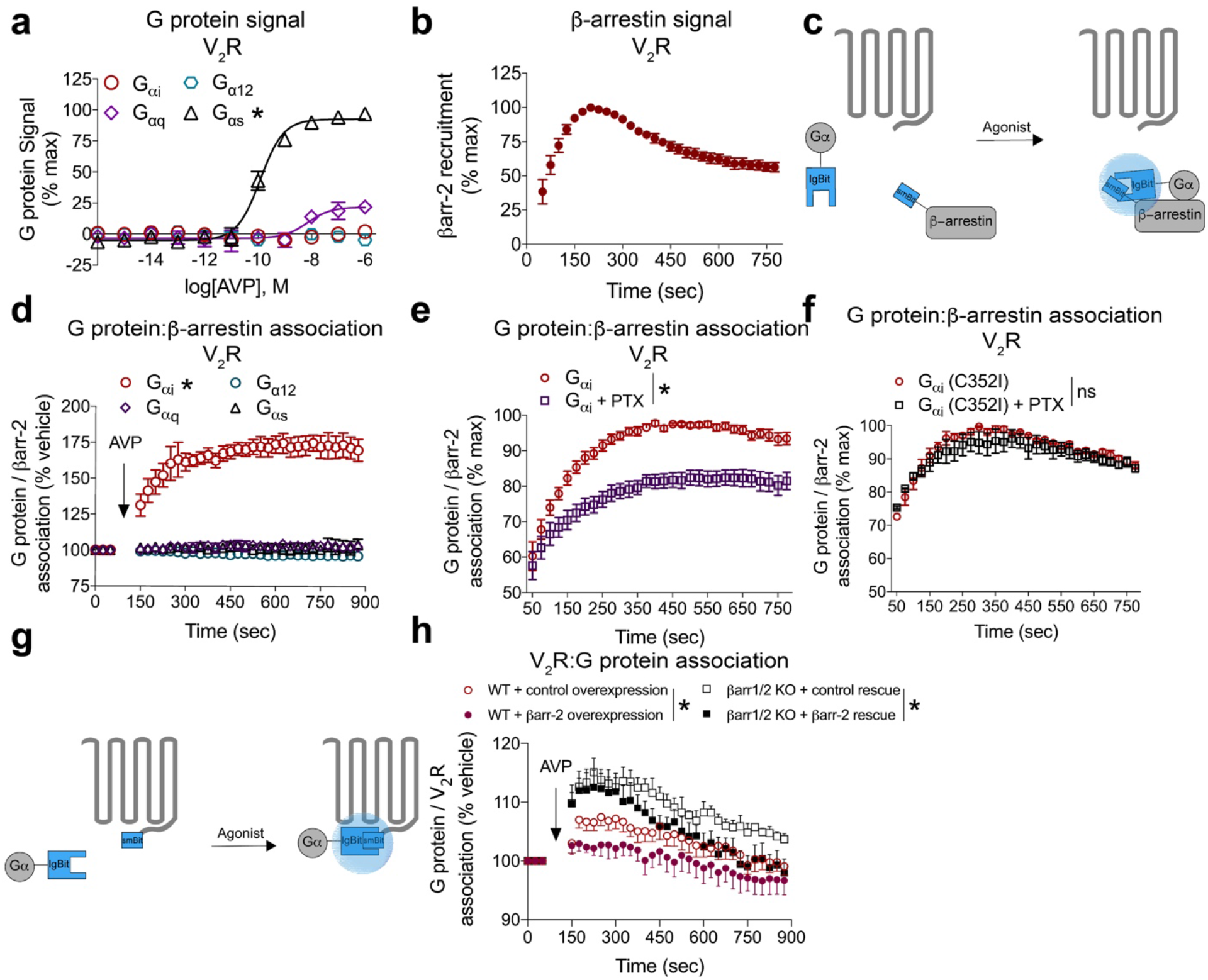
The canonically G_αs_-coupled V_2_R forms only G_αi_:β-arrestin complexes following AVP treatment. **a**, Assessment of canonical G protein signalling following agonist treatment of the V_2_R. **b**, Assessment of canonical β-arrestin-2 (smBiT) recruitment following agonist treatment of the V_2_R (LgBiT). **c**, Arrangement of luciferase fragments on G protein (LgBiT) and β-arrestin (SmBiT) in this two component assay. Unlike figures 1 and 2, the receptor is not tagged with a dipole acceptor. **d**, Effect of AVP (500 nM) treatment on cells overexpressing V_2_R in formation of Gα:β-arrestin-2 complexes. Only G_αi_ formed an observable complex with β-arrestin-2. **e**, Effect of pertussis toxin pretreatment on G_αi_:β-arrestin-2 complex formation. Data is normalized to maximal AVP-induced G_αi_:β-arrestin-2 signal within each replicate. **f**, Effect of pertussis toxin pretreatment on G_αi_ C352I mutant:β-arrestin-2 complex formation. **g**, Arrangement of luciferase fragments on G protein (LgBiT) and V_2_R (SmBiT) in this two component assay. **h**, Assessment of G_αi_ recruitment to the V_2_R following AVP treatment in either WT cells or β-arrestin-1/2 knockout cells and overexpressing or rescuing, respectively, with β-arrestin-2 or a pcDNA empty vector control. For panel **a**, experiments were conducted using the TGF alpha shedding assay in ‘ΔGsix’ HEK 293 cells. All other panels utilized WT HEK 293T cells overexpressing the indicated assay components. For panels **a** and **d**, \**P*<0.05 by two-way ANOVA, Fischer’s post hoc analysis with a significant difference G_αi_ subunit relative to all other G_α_ subunits. For panel h, *P*<0.05 by two-way ANOVA, main effect of β-arr-2 expression. For panels **e** and **f**, \**P*<0.05 by two-way ANOVA, main effect of pertussis toxin treatment. ns, not significant. Panels **a,b,d,f** n=3 per condition; for panel **h**, n=3-4; for panel **e** n=8. Graphs show mean ± s.e.m.

We proceeded to interrogate the formation of G protein and β-arrestin scaffolds using split luciferase technology^9^ (nanoBiT) by fusing the smaller subunit (smBiT) of the split luciferase to β-arrestin-2 and inserting the larger subunit (LgBiT) into a similar location in the four primary G_α_ families, G_αs_, G_αi_, G_αq_, and G_α12_. In direct contrast with the G_α_ protein signalling, only G_αi_, but not G_αs_, G_αq_ or G_α12_, formed an observable complex with β-arrestin following V_2_R treatment with AVP (Fig. 3d). Given that the V_2_R is not canonically known to signal through G_αi_, this was surprising, especially given the absence of G_αi_ signalling in our assay. Varying the amounts of G_α_ subunit transfected by up to 10-fold did not increase the interaction between non-G_αi_ family members and β-arrestin following agonist treatment of either the V_2_R or β_2_AR (Extended Data Fig. 6a-h). Furthermore, G_αi_ isoforms 2 and 3, as well as the highly homologous G_αo_, were all recruited to β-arrestin-2 with varying efficacy following agonist treatment of the V_2_R (Extended Data Fig. 7). A similar G_αi_-family interaction with β-arrestin-1 was also observed (Extended Data Fig. 8a,b).

The interaction between G_αi_ and β-arrestin was sensitive to *pertussis toxin* (Fig. 3e), which promotes enzymatic ADP ribosylation of cysteine 352 in helix 5 of G_αi_^14^. Mutation of cysteine 352 to isoleucine rescued the effect of *pertussis toxin* (Fig. 3f). Consistent with recent observations^13,15^, *pertussis toxin* pretreatment did not affect β-arrestin recruitment to either the V_2_R or β_2_AR (Extended Data Fig. 9a,b), which suggests that *pertussis toxin* did not reduce the efficacy of G_αi_:β-arrestin complex formation by interfering with β-arrestin recruitment to the receptor. In addition, β-arrestin was not necessary for the interaction of G_αi_ with the V_2_R, as previously validated HEK 293T cells lacking both β-arrestin-1 and β-arrestin-2 through CRISPR/Cas9 gene editing^16^ recruited G_αi_ following agonist treatment in both wild-type and β-arrestin1/2 knockout HEK 293T cell lines (Fig. 3g,h). Both β-arrestin-2 rescue in β-arrestin1/2 knockout cells or β-arrestin-2 overexpression in wild-type cells attenuated G_αi_:V_2_R association relative to the cell-type control (Fig. 3h). As expected, G_αs_ also associated with the V_2_R following agonist treatment (Extended Data Fig. 10). These results are consistent with findings that canonically G_αs_-coupled receptors can also recruit G_αi_^17^, without necessarily activating canonical G_αi_ signalling.

Given the paradoxical results of the G_αs_-coupled V_2_R catalyzing an unique G_αi_:β-arrestin complex, we proceeded to investigate if this phenomenon was generalizable to other GPCRs that canonically signal through different G_α_ proteins. We selected five well-studied GPCRs: the β_2_AR, CXCR3, NT_1_R, D_1_R, and D_2_R. Of these, only CXCR3 and D_2_R canonically signal through G_αi_. The β_2_AR and D_1_R canonically signal through G_αs_, and NT_1_R canonically signals through G_αq_. All five of these GPCRs formed G_αi_:β-arrestin complexes following agonist treatment (Figure 4).

**Figure 4:**
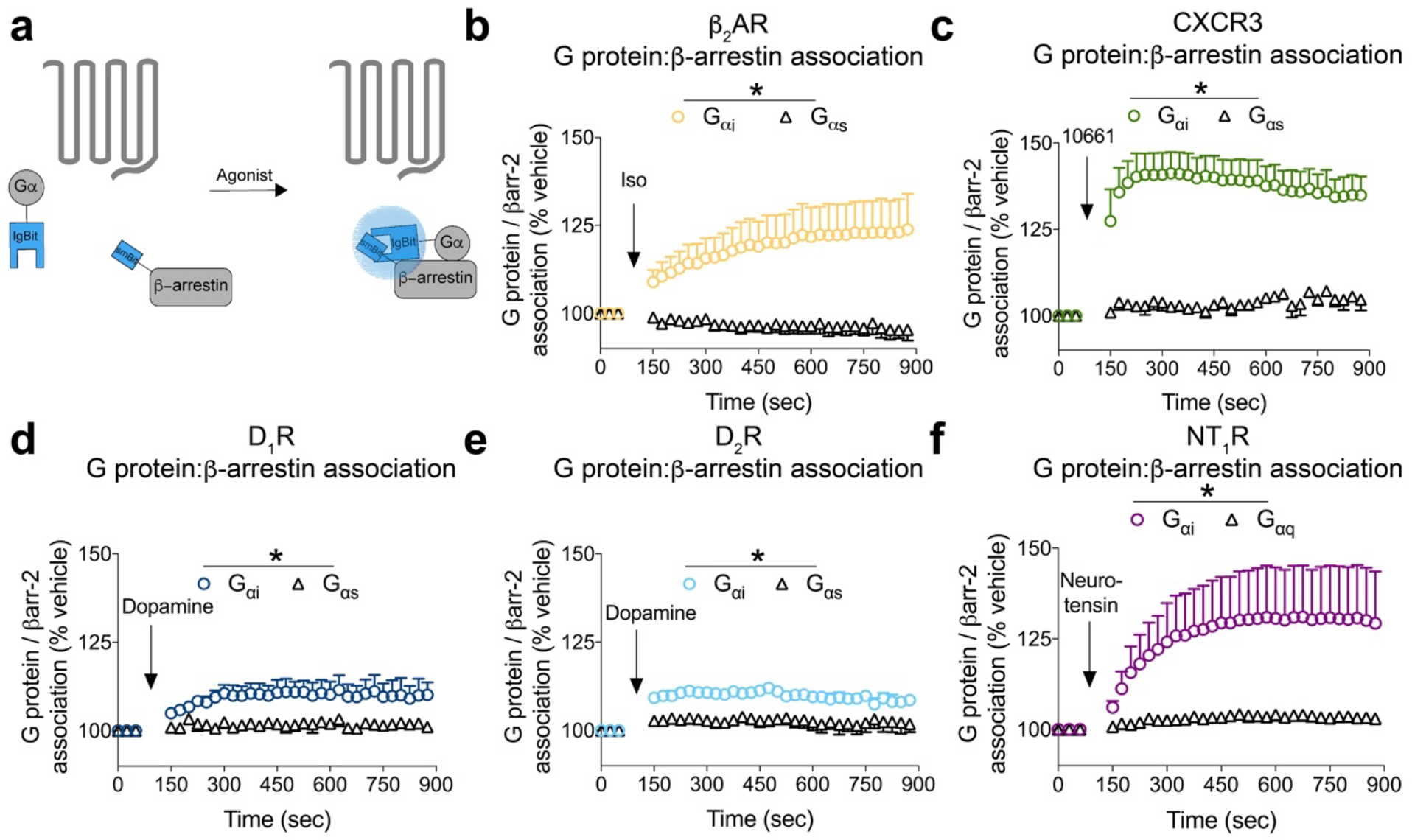
GPCRs form G_αi_:β-arrestin complexes following agonist treatment regardless of canonical G protein coupling. **a**, Arrangement of luciferase fragments on G protein (LgBiT) and β-arrestin (SmBiT) in this two component assay to assess the effect of the indicated agonist at forming G_αi_:β-arrestin-2 complexes in cells overexpressing **b**,β_2_AR (10 μM isoproterenol); **c**, CXCR_3_ (1 μM VUF10661); **d**, D_1_R (500 nM dopamine); **e**, D_2_R (500 nM dopamine); **f**, NT_1_R (10 nM neurotensin). \**P*<0.05 by two-way ANOVA, main effect of Gα subtype. For panel **b**, n=3-6; for panel **c**, n=3-4; for panel **d**, n=4, for panel **e**, n=3-4; for panel **f**, n=3 biological replicates per condition. Graphs show mean ± s.e.m.

### Complexes of G_αi_:β-arrestin facilitate ERK scaffolding and signalling

We next tested if the G_αi_:β-arrestin complex could scaffold ERK1/2 MAP kinases, which play critical roles in cell cycle regulation/proliferation and survival/apoptotic signalling^18^. A long held view is that GPCRs^19^ regulate ERK activation by inducing ERK phosphorylation through separate G protein and β-arrestin signalling pathways. Over a decade of work demonstrates that β-arrestins regulate ERK signalling, however, recent evidence using CRISPR/Cas9 genome editing approaches demonstrated a lack of ERK signalling in the collective absence of functional G proteins. This has led some to suggest that β-arrestin signalling is “dispensable” for ERK activation^20,21^. In apparent contrast, other studies have demonstrated an essential role for β-arrestin in some pathways regulating ERK activation^16^. Interestingly, two decades of work has demonstrated that for G_αi_ coupled receptors, β-arrestin-mediated ERK activation is invariably *pertussis toxin* sensitive^22,23,24^. However, how the G_αi_ and β-arrestin transducers cooperate in this pathway has remained obscure.

To investigate if G_αi_:β-arrestin complexes could directly scaffold to ERK downstream of the V_2_R, we utilized complex BRET by tagging ERK at either its N- or -C terminus with the dipole acceptor mKO (Fig. 5a). Data were normalized to an untagged (cytosolic) mKO to account for changes in protein localization following agonist treatment. Agonist treatment of V_2_R catalysed the formation of a G_αi_:β-arrestin:ERK complex (Fig. 5b). The magnitude of the adjusted complex BRET ratio was dependent on the location of the mKO tag on ERK (ERK-mKO compared to mKO-ERK), consistent with orientation and distance dependence of resonance energy transfer between luciferase donor and mKO dipole acceptor. To selectively evaluate the contributions of G_αi_ signalling on ERK phosphorylation, we utilized ‘ΔGsix’ HEK 293 cells depleted via CRISPR/Cas9 technology of G_αs_/G_αolf_, G_αq/11_, and G_α12/13_ G_α_ proteins previously reported and verified by western blot (Extended Data Fig. 11)^20^. Agonist treatment of these ‘ΔGsix’ HEK 293 cells overexpressing V_2_R robustly increased phosphorylated ERK (Fig. 5c,d). Pretreatment with *pertussis toxin* abrogated, but did not eliminate, ERK phosphorylation in ‘ΔGsix’ (Fig. 5c,d). Interestingly, in ‘ΔGsix’ cells, *pertussis toxin* pretreatment in combination with β-arrestin knockdown essentially eliminated ERK phosphorylation (Fig. 5c,d), consistent with functional coordination of G_αi_ and β-arrestin. These results would not have been predicted due to the known G_αs_-regulated ERK phosphorylation downstream of V_2_R and the demonstrated inability of V_2_R to canonically signal through G_αi_. However, this result is consistent with a role of G_αi_ coordination of β-arrestin signalling in this canonically G_αs_-coupled receptor.

**Figure 5:**
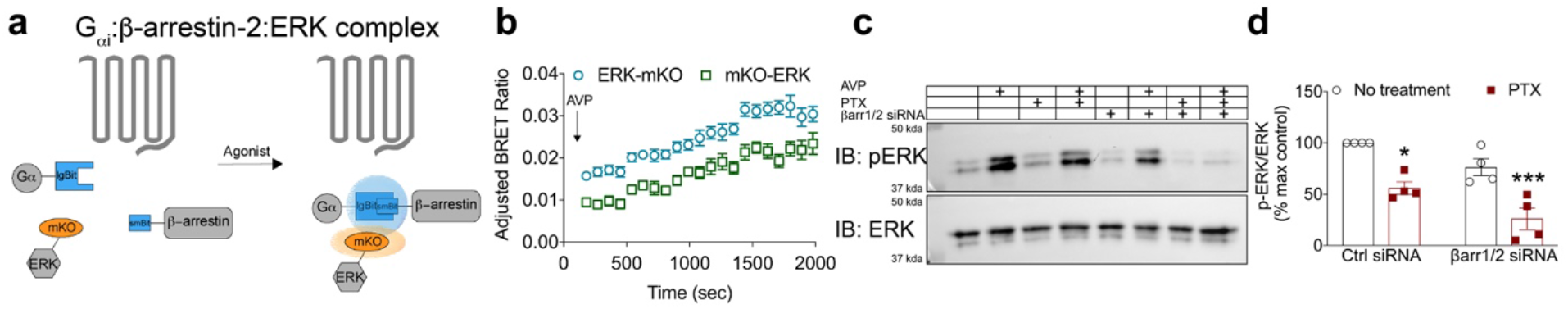
G_αi_:β-arrestin scaffolds form functional complexes with ERK. **a**, Arrangement of luciferase fragments and mKO acceptor fluorophore for complex BRET on G protein (LgBiT), β-arrestin (SmBiT), or ERK2 (mKO). **b**, Complex BRET association of G_αi_, β-arrestin, and ERK2 in cells overexpressing untagged V_2_R following treatment with AVP (500 nM). Data were normalized to both vehicle treatment and cytosolic mKO. **c**, Representative immunoblot of phospho and total ERK1/2 in ‘ΔGsix’ HEK 293 cells pretreated with PTX (200 ng/mL) and/or βarr1/2 siRNA, stimulated with either vehicle or AVP (500 nM). ERK1/2 phosphorylation was nearly eliminated in cells treated with both PTX and βarr1/2 siRNA. **d**, Quantification of ERK immunoblots. \**P*< 0.05, \*\*\**P*< 0.001, two-way ANOVA with Bonferroni post hoc to no treatment, control siRNA group. The net BRET ratio of cytosolic mKO control was subtracted from the net BRET ratio of ERK-mKO to yield an adjusted BRET ratio that is the ordinate of panel **b**. Immunoblots are representative of three separate experiments. For panel **b**, n=5; panel **d**, n=4. Immunoblot is representative of four experiments. PTX, pertussis toxin. Graphs show mean ± s.e.m.

### A β-arrestin-biased agonist promotes formation of the G_αi_:β-arrestin complex and displays *pertussis* toxin-sensitive cell migration

As described above it has long been clear that G_αi_ and β-arrestin somehow coordinate their signalling to ERK downstream of canonically G_αi_ coupled receptors, whereas in the case of GPCRs coupled to other G proteins these two signalling arms have appeared more independent^25^. However, our observation that this G_αi_:β-arrestin complex forms downstream of receptors not typically thought to interact with G_αi_ suggests that the formation of G_αi_:β-arrestin complexes may be widespread. To further assess this model we utilized the angiotensin type 1 receptor (AT_1_R) β-arrestin-biased agonist, TRV120023, which is well-established to have no appreciable canonical G protein signalling but robustly recruits β-arrestin to the AT_1_R^26,27^ (Fig. 6a). This contrasts with the endogenous ligand of AT_1_R, Angiotensin II (AngII), which when applied to the AT_1_R signals through both G_αq_ and G_αi_ (Fig. 6b), as well as recruits β-arrestin (Fig 6a). We verified that TRV120023 is a β-arrestin-biased agonist in our assays and that it had no appreciable ability to promote canonical G protein signalling through any of the four G_α_-family proteins tested (Fig. 6c) while strongly stimulating β-arrestin recruitment to the receptor (Fig. 6a). Because TRV120023 does not appreciably activate canonical G protein signalling, it would be predicted that it would not induce cell migration, a function thought to require canonical G protein signalling. However, not only did TRV120023 promote cellular migration, this migration was *pertussis toxin* sensitive, as pretreatment of cells with *pertussis toxin* reduced TRV120023-mediated migration by ~50%. Furthermore, inhibition of both G_αi_ and β-arrestin through *pertussis toxin* pretreatment and siRNA knockdown of β-arrestin1/2 eliminated migration (Fig. 6d,e). Similar to all other receptors tested in the current study, both the endogenous agonist AngII and the β-arrestin-biased agonist TRV120023 induced G_αi_:β-arrestin complex formation (Figure 6f).

**Figure 6:**
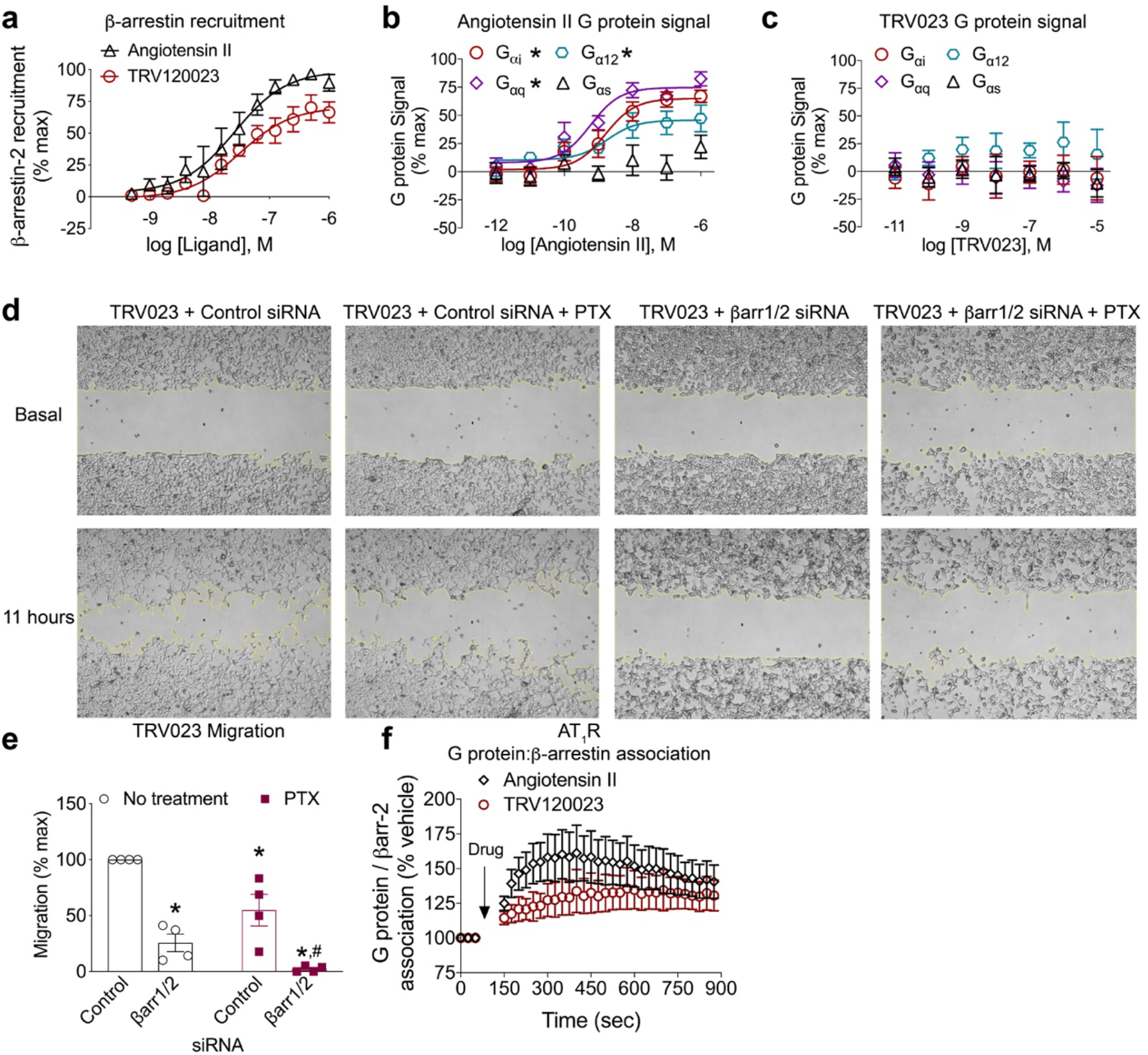
Cell migration to the β-arrestin-biased Angiotensin ligand TRV120023 requires both G_αi_ and β-arrestins. **a**, BRET assay quantifying the recruitment of β-arrestin-2-YFP to AT_I_R-RlucII following treatment with either angiotensin II or TRV120023. Assessment of canonical G protein signalling at the Angiotensin II type 1 receptor (AT_I_R) following treatment with either the endogenous ligand, angiotensin II (**b**) or the previously characterised β-arrestin-biased ligand TRV120023 (**c**). **d**, Representative images of the four TRV120023 migration conditions in HEK 293 cells stably expressing AT_I_R. **e**, Quantification of PTX pretreatment (200 ng/mL) and/or β-arr1/2 siRNA on TRV120023-induced migration for the experiment shown in panel **d. f**, Split luciferase assay for monitoring G protein-β-arrestin association after treatment of AT_I_R with either angiotensin II or TRV120023. *P< 0.05, two-way ANOVA with Bonferroni post hoc to no treatment, control siRNA group. #P< 0.05, two-way ANOVA with Bonferroni post hoc compared to control siRNA, PTX pretreated group. For panel **a**, n=4 per condition, for panels **b** and **c**, n=4-5 per condition, for panels **e** and **f**, n=4 per condition. Graphs show mean ± s.e.m.

## Discussion

Our results reveal a new GPCR signalling paradigm in which GPCRs can promote formation of a G_αi_:β-arrestin complex. The formation of this G_αi_:β-arrestin complex was observed downstream of all receptors tested, even those receptors that do not canonically signal through G_αi_. A unique feature of our findings is the ability of a variety of GPCR ligands to drive formation of G_αi_:β-arrestin complexes, even a β-arrestin-biased ligand that has little or no ability to promote G protein-mediated signalling. This suggests that a major driver of the association of β-arrestin with G_αi_ is the GPCR-mediated recruitment of β-arrestin to the plasma membrane. The observed G_αi_:β-arrestin scaffolds can include a GPCR or a signalling effector (ERK), or possibly both, and suggest that G_αi_:β-arrestin scaffolds form functional signalling complexes. Remarkably, these signalling complexes are associated with ERK activation, even when the stimulatory GPCR ligand is incapable of activating canonical G_αi_ signalling. Using HEK293 cells depleted of the G_αs/q/12_ proteins and overexpressing the V_2_R, we demonstrate that AVP-induced ERK phosphorylation is nearly eliminated following G_αi_ inhibition with *pertussis toxin* and siRNA knockdown of β-arrestins. Consistent with these results, we show that *pertussis toxin* impairs migration of cells treated with a β-arrestin-biased ligand, TRV120023. While these results are concordant with functional G_αi_:β-arrestin scaffolds, it remains unclear how G_αi_:β-arrestin complexes participate in the process of ERK activation and cell migration.

This study bridges seemingly contradictory results concerning the interplay of G protein and β-arrestin signalling^16,20,21^ by delineating a novel G_αi_:β-arrestin scaffolding complex. A number of significant caveats must be considered when interpreting our results. Most importantly, these studies rely on the overexpression of components which have been genetically modified by insertion of various fluorescent reporter probes. For example, as demonstrated in Figure 1, a particular probe architecture can have significant impact on the intensity of the signal generated. Moreover, as a consequence of overexpression, interactions may be detected which would not be seen at physiologically relevant concentrations of these molecules. However, our control experiments, and our observation that the *lowest* concentrations of expression vector provide the *highest* signal-to-noise for G_αi_:β-arrestin complex formation (Extended Data Fig. 6) support our interpretation that this complex formation is neither an artifact of probe orientation nor enhanced by protein overexpression. The presence of G_αi_:β-arrestin association in orthogonal assays (TSA and coimmunoprecipitation) provides further support for the existence of G_αi_:β-arrestin complexes. These experiments offer plausible mechanistic insight into initially paradoxical observations that G_αi_ can drive ERK phosphorylation downstream of the canonically G_αs_-coupled V_2_R and that *pertussis toxin* inhibits cell migration to a β-arrestin-biased agonist. Further studies examining both the biochemical mechanisms underlying, as well as additional functions of G_αi_:β-arrestin scaffolds, will be required to address their physiological role and therapeutic implications.

## Methods

### Cell culture and transfection

Human embryonic kidney cells (HEK 293, HEK 293T, Rockman βarrestin-1/2 HEK 293 knockout, and ‘ΔGsix’ HEK 293) were maintained in minimum essential medium supplemented with 1% anti-anti and 10% fetal bovine serum. Rockman βarrestin-1/2 HEK 293 knockout were supplied by Dr. Howard Rockman and validated as previously described^16^. Cells were grown at 37 °C with humidified atmosphere of 5% CO_2_. For BRET and luminescence studies, HEK 293T cells were transiently transfected via an optimized calcium phosphate protocol as previously described. For immunoblot studies utilizing siRNA, HEK 293T cells were transiently transfected with Lipofectamine 3000 (ThermoFisher) according to manufacturer specifications. For TGF alpha shedding assay studies, ‘ΔGsix’ HEK 293 cells were transfected using Fugene 6 (Promega) according to manufacturer specifications.

### Generation of constructs

Cloning of constructs was performed using conventional techniques such as restriction enzyme/ligation methods. Linkers between the fluorescent proteins or luciferases and the cDNAs for receptors, transducers, kinases, or adaptor proteins were flexible (GGGGS) and ranged between 15-18 amino acids. See supplementary table for complete list of constructs used in manuscript.

### Split luciferase and complex BRET assays

HEK293T cells seeded in 6-well plates were co-transfected with 500 ng of smBiT tagged β-arrestin-2, and either 250 ng of LgBiT tagged receptor or 2000 ng of untagged receptor and varying concentrations of LgBiT G_α_ protein expression vector (most experiments were conducted between 50-200 ng of G_α_ plasmid) or 2000ng of mKO tagged β-arrestin-2 and 500 ng of smBiT tagged V_2_R using a calcium phosphate protocol previously described^28^. Twenty-four hours post-transfection, cells were plated onto clear bottom, white-walled 96-well plates at 50,000-100,000 cells/well in “BRET media” - clear minimum essential medium (GIBCO) supplemented with 2% FBS, 10 mM HEPES, 1x GlutaMax, and 1x Anti-Anti (GIBCO). Select cells were then treated overnight with *pertussis toxin* pretreatment at a final concentration of 200 ng/mL. The following day, media were removed, and cells were incubated at room temperature with 80 μL of Hanks’ balanced salt solution (GIBCO) supplemented with 20mM HEPES and 3 μM coelenterazine-h for 15 minutes. For luminescence split luciferase studies, plates were read with a BioTek Synergy Neo2 plate reader set at 37 °C with a 485 nm emission filter. Cells were stimulated with either vehicle (Hank’s Balanced Salt Solution with 20 mM HEPES) or indicated concentration of agonist. For split luciferase luminescence experiments, plates were read both before and after ligand treatment to calculate Δnet change in luminescence and subsequently normalized to vehicle treatment. For complex BRET experiments, plates were read on a Berthold Mithras LB940 using pre-warmed media and instrument at 37 °C using a standard Rluc emissions filter (480 nm) with a custom mKO 542 nm long-pass emission filter (Chroma Technology Co., Bellows Falls, VT). Readings were performed using a kinetic protocol with automatic injection of ligands as indicated in figures. The BRET ratio was calculated by dividing the mKO signal by the luciferase signal. For some experiments, a Net BRET ratio was calculated by subtracting the vehicle BRET ratio from the ligand stimulated BRET ratio, or an adjusted BRET ratio was calculated by subtracting the ligand treated cytosolic mKO signal from the ligand treated effector mKO signal, as indicated in figure legends.

### Immunoblotting

Experiments were conducted as previously described^28^. Briefly, cells were serum starved for at least four hours, stimulated with the indicated ligand, subsequently washed 1x with ice-cold PBS, lysed in ice-cold RIPA buffer containing phosphatase and protease inhibitors (Phos-STOP (Roche), cOmplete EDTA free (Sigma)) and rotated for forty-five minutes, and cleared of insoluble debris by centrifugation at >12,000 x g (4 °C, 15 minutes), after which the supernatant was collected. Protein was resolved on SDS-10% polyacrylamide gels, transferred to nitrocellulose membranes, and immunoblotted with the indicated primary antibody overnight (4°C). phospho-ERK (Cell Signaling Technology, #9106) and total ERK (Millipore #06-182) were used to assess ERK activation. A1-CT antibody that recognizes both isoforms of β-arrestin was utilized^16^, with protein loading assessed by alpha-tubulin (Sigma #T6074). Galpha i-1 (13533, Santa Cruz Biotechnology), Galpha q/11/14 (365906, Santa Cruz Biotechnology), Galpha 12 (515445, Santa Cruz Biotechnology), Galpha 13 (293424, Santa Cruz Biotechnology), Galpha s/olf (55545, Santa Cruz Biotechnology) antibodies were used to verify ‘ΔGsix’ HEK 293 cells. Horseradish peroxidase-conjugated polyclonal mouse anti-rabbit-IgG or anti-mouse-IgG were used as secondary antibodies. Immune complexes on nitrocellulose membrane were imaged by SuperSignal enhanced chemiluminescent substrate (Thermo Fisher). Following detection of phospho signal, nitrocellulose membranes were stripped and reblotted for total kinase signal. For quantification, phospho-protein signal was normalized to total protein signal using ImageLab (Bio-Rad) within the same blot. siRNA knockdown of β-arrestin-1 and β-arrestin-2 was conducted as previously described.

### siRNA knockdown

HEK 293T cells were transiently transfected with Lipofectamine 3000 (Thermo Fisher) per manufacturer specifications in a six-well tissue culture sterile plate with 1 ug of receptor and 3.5μg of either control or siRNA directed to β-arrestin-1/2 sequences (“wem2”) as previously described^16^.

### Wound-healing migration assay

HEK 293T cells stably expressing the AT_1_R were utilized. Briefly, 70 μl of cell suspension at a concentration of 5 x 10^5^ cells per mL was applied into each well of silicone inserts (Ibidi, Martinsried, Germany) on 24 well plate, and after 24 hrs incubation, the inserts were removed to create a wound field. The cells were incubated additionally for 12 hrs with 1 μM of TRV120023 and visualized with a Zeiss Axio Observer microscope (Carl Zeiss, Thornwood, NY). Wound healing was then analysed using ImageJ (NIH, Bethesda, MD) wound healing tool macros.

### TGF-alpha shedding assay

GPCR G_α_ activity was assessed by the transforming growth factor-α (TGF-α) shedding assay as previously described^29^. Briefly, HEK 293 cells lacking G_αq_, G_α11_, G_αs/olf_, and G_α12/13_ (‘ΔGsix’ HEK 293 cells) were transiently transfected with receptor, modified TGF-α-containing alkaline phosphatase (AP-TGF-α), and the indicated G_α_ subunit. Cells were reseeded twenty-four hours later in Hanks’ Balanced Salt Solution (HBSS) (Gibco, Gaithersburg, MD) supplemented with 5mM HEPES in a Costar 96-well plate (Corning Inc., Corning, NY). Cells were then stimulated with the indicated concentration of ligand for one hour. Conditioned media (CM) containing the shed AP-TGF-α was transferred to a new 96-well plate. Both the cell and CM plates were treated with para-nitrophenylphosphate (p-NPP, 100mM) (Sigma-Aldrich, St. Louis, MO) substrate for one hour, which is converted to para-nitrophenol (p-NP) by AP-TGF-α. This activity was measured at OD_405_ in a Synergy Neo2 Hybrid Multi-Mode (BioTek, Winooski, VT) plate reader immediately after p-NPP addition and after one-hour incubation. G_α_ activity was calculated by first determining p-NP amounts by absorbance through the following equation: 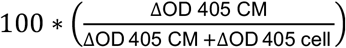 where ΔOD 405 = OD 405 1 hour – OD 405 0 hour and ΔOD 405 cell and ΔOD 405 CM represent the changes in absorbance after one hour in the cell and CM plates, respectively. Data were normalized to a single well that produced the maximal signal.

### Thermal Shift Assay

Protein thermal melting shift experiments were performed using the StepOnePlus™ Real-Time PCR System (Applied Biosystems). Proteins were buffered in 20 mM HEPES pH 7.5, 100 mM NaCl, 4 mM MgCl_2_. β-arrestin-2, G_αiβγ_, V_2_Rpp, Fab30, and nonhydrolyzable GTP analog of GTP GMP-PNP were added at a final concentration of 5 μM, 10 μM, 30 μM, and 120 μM, respectively. All reactions were set up in a 96-well plate at final volumes of 20 μl and SYPRO Orange (Thermo Fisher Scientific) was added as a probe at a dilution of 1:1000. Excitation and emission filters for the SYPRO-Orange dye were set to 475 nm and 580 nm, respectively. The temperature was raised with a step of 0.5 °C per 30 second from 25 °C to 99 °C and fluorescence readings were taken at each interval. All measurements were carried out three times. Data were analysed using Applied Biosystems^®^ Protein Thermal Shift™ Software. Expression and purification of heterotrimeric G protein was conducted as previously described^30^. In brief, Hive Five insect cells were infected with two viruses made from BestBac baculovirus system, one expressing human Gβ_1_-His6 and G_γ2_ and the other G_αi1_. Approximately forty-eight hours after infection the cells were harvested, solubilized, and heterotrimeric G_αi_ purified using Ni-NTA chromatography and HiTrap Q sepharose anion exchange (GE Healthcare Life Sciences).

### Immunoprecipitation

Immunoprecipitation was conducted as previously described^31^. Briefly, 4 μg of HA-V_2_R, 4 μg of G_αi_-GFP and 4 μg of pcDNA-ARRB1-Flag and/or pcDNA were transfected into HEK 293 cells seeded in 6 cm plates. Forty-eight hours post-transfection, after approximately 4 hours of serum starvation, cells were stimulated with AVP for 5 and 10 mins at 37 °C. Cells were then lysed on ice for 10 min in FLAG lysis buffer (50 mM Tris-HCl, pH 7.4, 1% Triton X-100, 150 mM NaCl, 1 mM EDTA) supplemented with protease inhibitor cocktail tablet (Roche). Cell lysates were incubated with anti-FLAG M2 affinity gel (A2220, Sigma) overnight and immunoprecipitated ARRB1-FLAG were eluted with Flag peptides (F3290, Sigma). For primary antibody incubation, GFP polyclonal antibody (A6455, Invitrogen), HA-Tag (3724S, cell signaling biotechnology), and ANTI-FLAG M2 antibody (F3165, Sigma) were utilized.

### Confocal Microscopy

HEK293T cells plated in fibronectin-coated 35 mm glass bottomed dishes (MatTek Corp. P35G-0-14-C) were transiently transfected via the calcium-phosphate method with DNA encoding G_αi_-mVenus (125 ng), βarr2-mKO (125 ng), and/or Mars1-V_2_R (500 ng). Mars1 binds a membrane impermeant fluorogen (SCi1) and induces its fluorescence in the near-infrared spectrum^32^. Cells were pulse labelled with SCi1 (diluted 1:5000 from 0.5 mg/mL stock) for 15 minutes before treatment with or without 100 nM AVP. Cells were then fixed at basal, 5 minutes and 30 minutes after treatment with 4% paraformaldehyde. Samples were then imaged with a Plan-Apochromat 63x/1.4 Oil lens on a Zeiss LSM880 using corresponding laser lines to excite mVenus, mKO, or Mars1 (488*nm*, 561 *nm*, 633*nm* respectively). Spectral gating via a 34 spectral array detector was performed using single colour transfection controls.

### Drugs

VUF10661, AVP, dopamine, angiotensin II, neurotensin and isoproterenol were all purchased from Sigma-Aldrich (St. Louis, MO). VUF10661 and isoproterenol were dissolved in dimethyl sulfoxide (DMSO) to make stock solutions and stored in a desiccator cabinet. Stock solutions of AVP, angiotensin II, neurotensin, (Sigma-Aldrich) were prepared according to manufacturer specifications. TRV120023 was provided by Trevena (King of Prussia, PA). Stock solutions of neurotensin were made in 0.1% BSA in PBS. Dopamine was prepared fresh in BRET media supplemented with 0.03% ascorbic acid (Sigma-Aldrich). All drug dilutions were performed with BRET media or cell culture media. PTX was obtained from List Biological Laboratories (Campbell, CA). All compound stocks were stored at −20°C until use.

### Data availability

The data sets generated for this study are available from the corresponding author upon reasonable request. All relevant data are included in the paper or the supplementary information.

### Statistical analyses

Dose-response curves were fitted to a log agonist versus stimulus with three parameters (span, baseline, and EC50) with the minimum baseline corrected to zero using Prism 8.0 (GraphPad, San Diego, CA). For comparing ligands in concentration-response assays or time-response assays, a two-way ANOVA of ligand and concentration or ligand and time, respectively, was conducted. Unless otherwise noted, statistical tests were two-sided and corrected for multiple comparisons. Further details of statistical analysis and replicates are included in the figure captions.

**Table 1:**
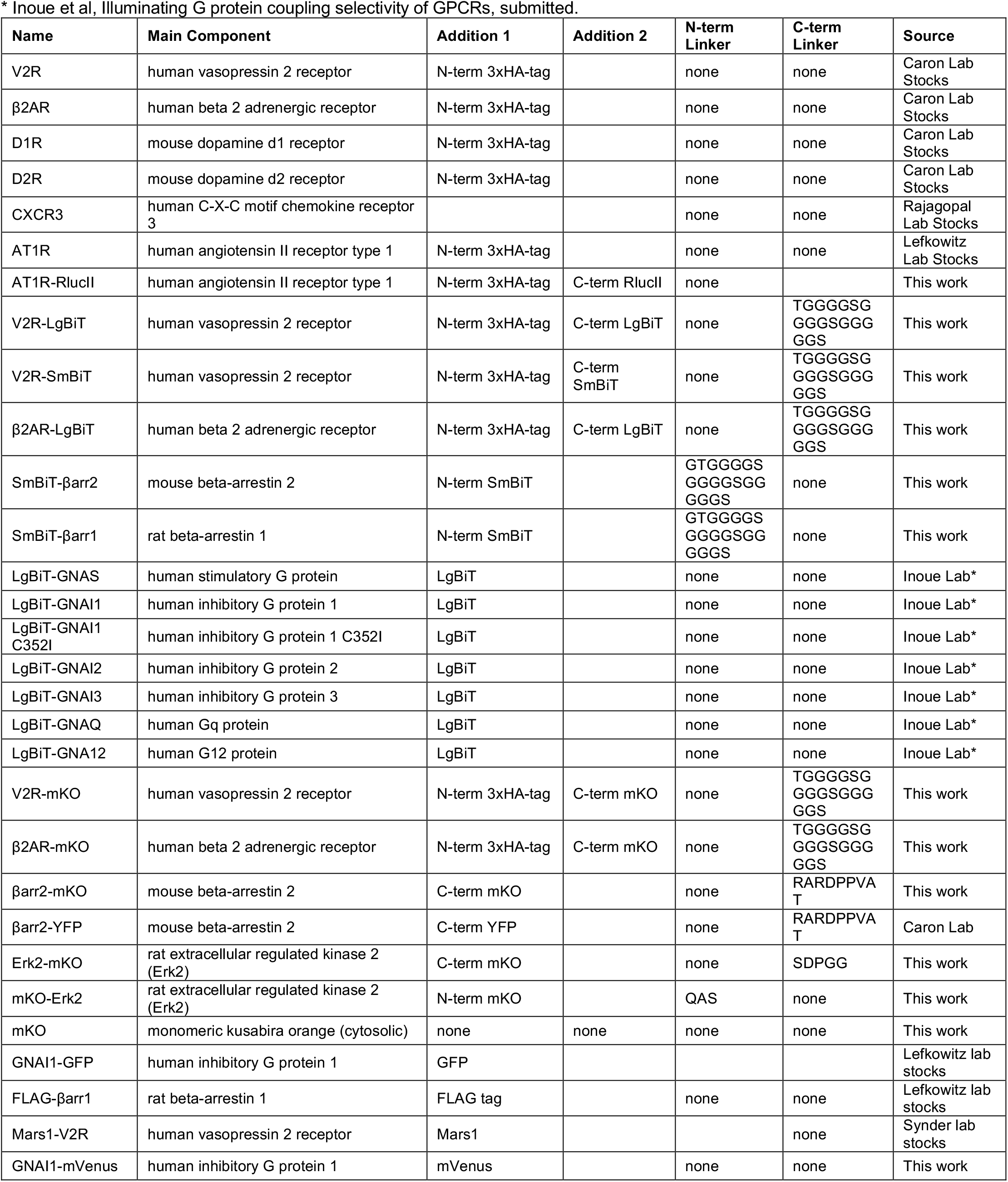
Constructs Used.

## Acknowledgements

The authors recognize consequential contributions from R. J. Lefkowitz for his helpful comments and suggestions regarding experimental design, laboratory resources (including constructs outlined in the methods section), data interpretation, and edits to the text throughout multiple drafts. The authors thank N. Nazo for administrative assistance; L. Luttrell, S. Shenoy, C. Chavkin, G. Viswanathan and J. Silverman for helpful discussion and thoughtful feedback; M. Orlen and D. Eiger for technical assistance; S. Shenoy and N. Freedman for the use of laboratory equipment, and Dr. H Rockman for Rockman β-arrestin-1/2 KO HEK293 cells. This work was supported by T32GM7171 (J.S.S.), the Duke Medical Scientist Training Program (J.S.S.), F31DA041160 (T.F.P.), PRIME JP17gm5910013 (A.I.), the LEAP JP17gm0010004 from the Japan Agency for Medical Research and Development (A.I.), the JSPS KAKENHI (A.I.), 17K08264R37MH073853 (M.G.C.), 1R01GM122798-01A1 (S.R.), K08HL114643-01A1, (S.R.), Burroughs Wellcome Career Award for Medical Scientists (S.R.).

## Author Contributions

J.S.S. and T.F.P. contributed equally to this work. J.S.S. and T.F.P conceived of the study and designed, generated, and validated receptor and β-arrestin split luciferase and mKO constructs. A.I. designed, generated, and validated all G protein split luciferase constructs and generated ‘ΔGsix’ cells. J.S.S., T.F.P., C.L., K.Z., I.C., X.X., Z.M., I.M.L performed cell-based experiments. X.X. performed the migration assay and analysed the data. A.W.K. performed and analysed TSA experiments. D.P.S. contributed purified protein for TSA experiments. L.K.R and J.C.S performed and analysed confocal experiments. J.S.S., T.F.P., and S.R. analysed all other data. J.S.S., T.F.P., M.G.C, and S.R. wrote the paper. All authors discussed the results and commented on the manuscript.

---

**Extended Data Figure 1:**
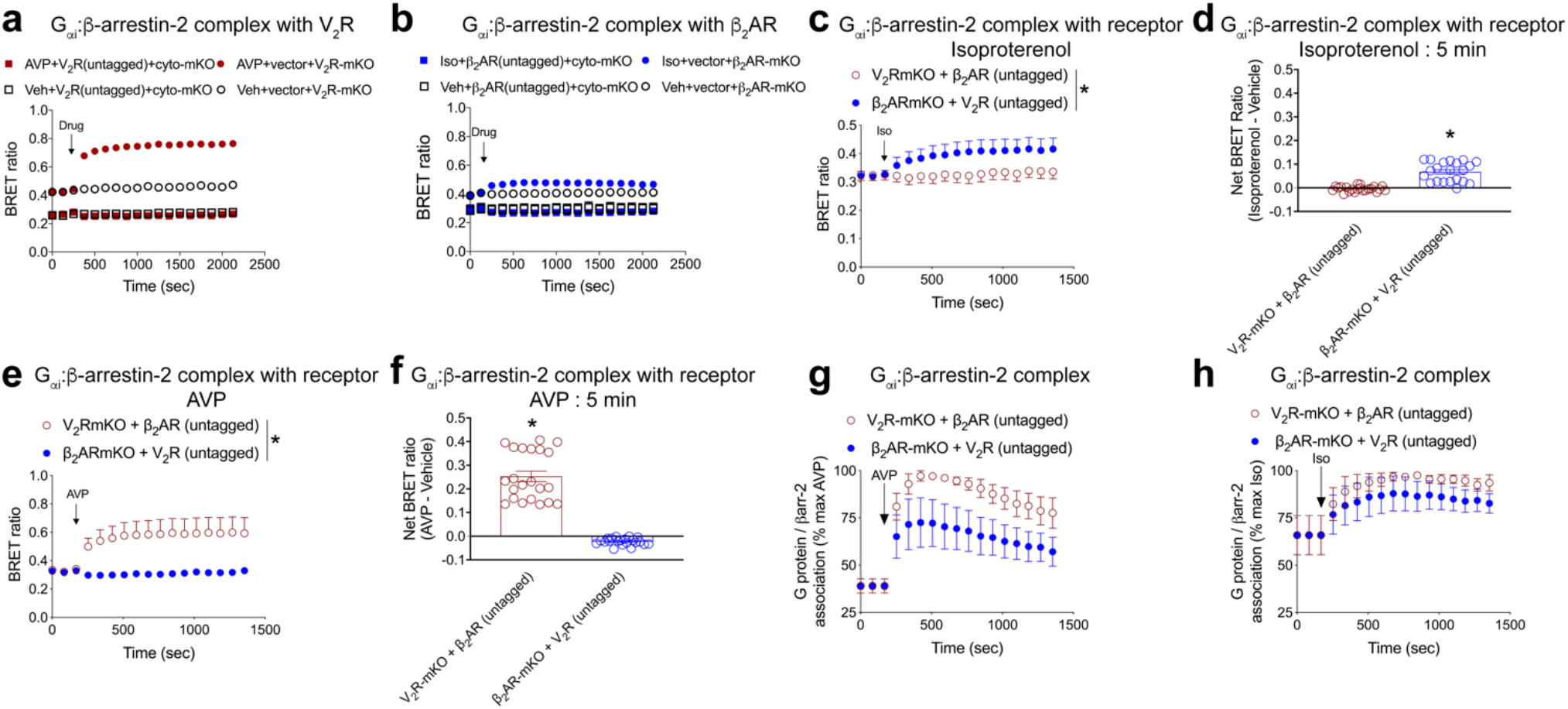
Additional complex BRET controls. HEK 293T cells were transiently transfected with the indicated receptor(s) and assay components. Cytosolic mKO (untagged) was utilized as a non-specific dipole acceptor control. **a**, G protein-β-arrestin complex association with mKO. mKO was expressed either in the cytosol (untagged) or tagged to the C-terminus of V_2_R. For the cytosolic mKO groups, untagged V_2_R was transfected to allow for AVP-induced G protein:β-arrestin association. To kinetically assess ligand-induced increases in V_2_R:G protein:β-arrestin formation, three baseline reads were conducted, followed by treatment with either vehicle or AVP (500 nM). Consistent with a selective interaction, only AVP treatment of the V_2_R-mKO condition resulted in formation of a V_2_R:G_αi_ protein:β-arrestin megaplex. Baseline differences in the BRET ratio observed between cytosolic mKO and V_2_R-mKO most likely reflect differences in mKO localization with in the cell. **b**, Similar experiment to panel **a**, except assessing a β_2_AR G_αi_:β-arrestin megaplex following treatment with either vehicle or isoproterenol (10 μM) after three baseline reads. mKO was either expressed in the cytosol or tagged on the C-terminus of β_2_AR. Panels **c-h**, experiments were conducted to test the specificity of the complex BRET assay to test if an untagged receptor stimulated with its cognate ligand could form a ‘bystander,’ non-specific megaplex. In panels **c-h**, replicate experiments were conducted within the same plates under the indicated assay conditions to minimize plate to plate variation. **c**, Only when AVP is paired with V_2_R-mKO is a V_2_R:G_αi_:β-arrestin megaplex formed, and no increase in the BRET ratio is observed under conditions with cells expressing β_2_AR-mKO and untagged V_2_R. This indicates complex BRET selectively measures GPCR megaplexes and minimizes bystander effects. **d**, 5 minute time point of data shown in panel c. **e**, Similar experiment to panel c, except assessing the ability of isoproterenol (10 μM) to quantify β_2_AR:G_αi_:β-arrestin in cells expressed either native β_2_AR with V_2_R-mKO or β_2_AR-mKO with untagged V_2_R. Similar to panel c, only when isoproterenol is paired with β_2_AR-mKO is a β_2_AR:G_αi_:β-arrestin megaplex formed, and no increase in the BRET ratio is observed under conditions with cells expressing β_2_AR-mKO and untagged V_2_R. **f**, 5 minute time point of data shown in panel **e**. In panels **g** and **h**, only luciferase complementation in the 480nm channel that indicates G_αi_:β-arrestin complex formation is shown (no BRET data is included). The same cells and conditions utilized in panels **c-f** are used. This control experiment assessed the ability of either native V_2_R or V_2_R-mKO to induce association of G protein-β-arrestin. **g**, As expected from data shown in Figures 3 and 4 in the main text, AVP (500 nm) induced association of G_αi_ and β-arrestin-2 in cells expressing either native V_2_R or V_2_R-mKO. Slight deviations in efficacy likely reflect minor differences in surface expression. **h**, Similar experiment to panel **g**, except assessing the ability of isoproterenol (10 μM) to induce association of G_αi_:β-arrestin in cells expressing either untagged β_2_AR or β_2_AR-mKO. Similar to panel **g**, slight deviations in efficacy likely reflect minor differences in receptor surface expression. For panels **c, e**, \**P*< 0.05, two way ANOVA with main effect of construct. For panels **d, f**, \**P*< 0.05, two-tailed t-test. Panels **a,b**, n=4 per condition; panels **c-h**, n=3 per condition. Individual wells in the 96 well plates of the 3 different replicates are shown in panels **d** and **f** for the purpose of displaying experimental variability. Graphs show mean ± s.e.m.

**Extended Data Figure 2:**
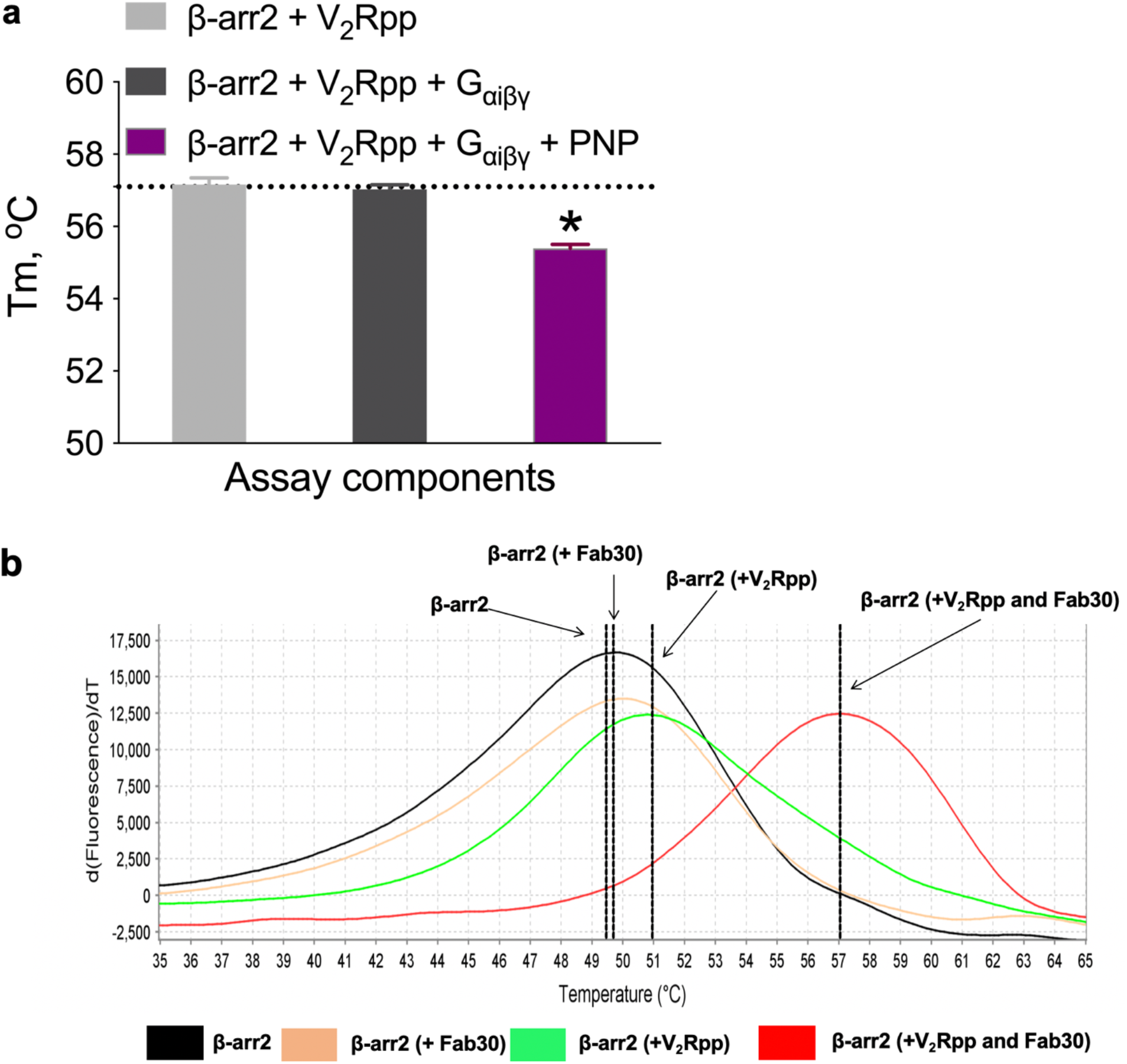
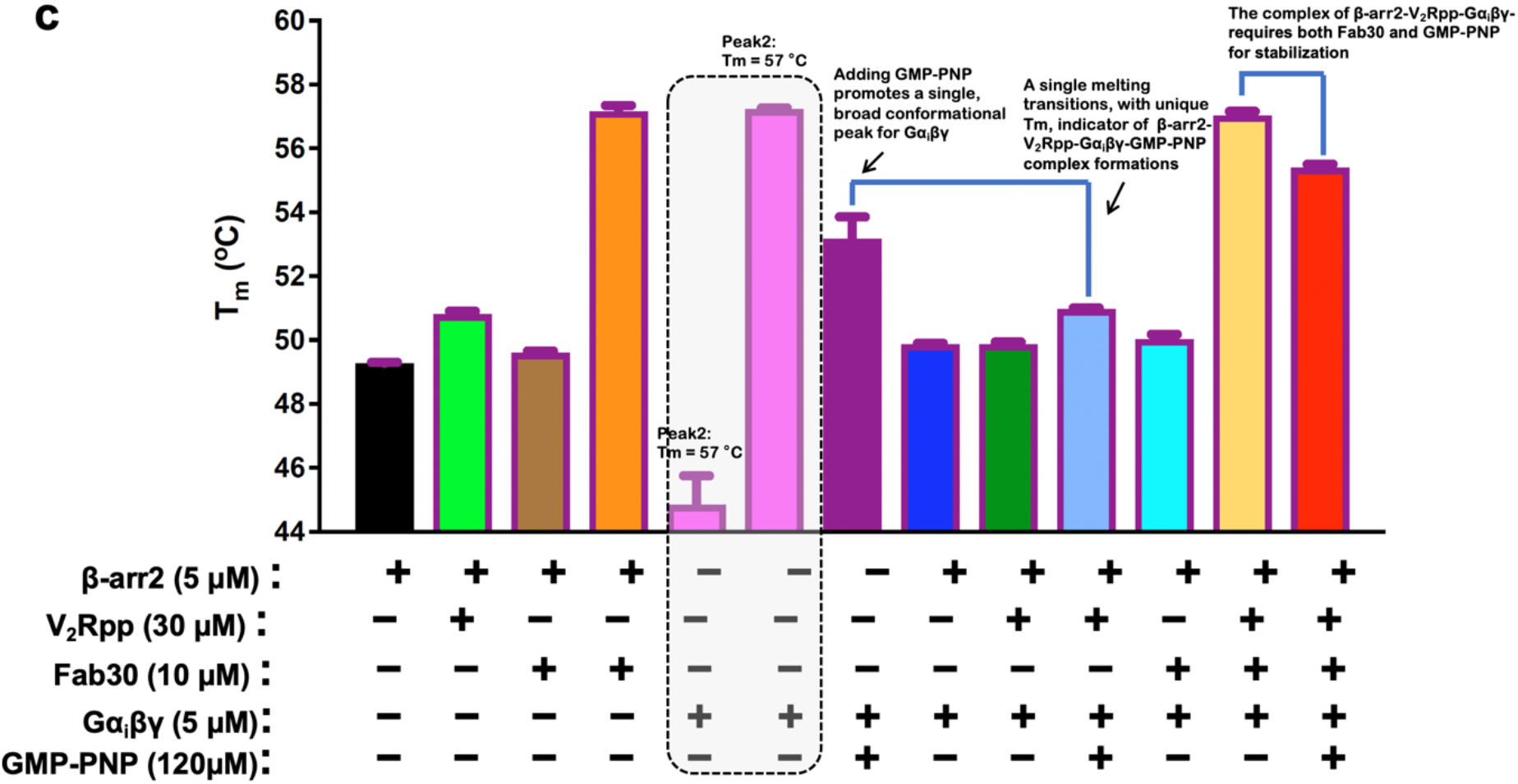
Thermal stability assay melting curves. **a**, Thermal stability assay of purified complex components including β-arrestin-2, phosphorylated vasopressin C-terminal peptide (V_2_Rpp), with heterotrimeric G_αiβγ_ ± non-hydrolyzable GTP (PNP). All TSA experiments contained the stabilizing antigen binding fragment 30 (Fab30). The change in melting temperature with the indicated assay components is consistent with complex formation. **b**, melt profiles of β-arrestin-2 alone (black), in presence of a GPCR V_2_-receptor C-terminal tail phosphopeptide (V_2_Rpp) (green), Fab30 (orange) or V_2_Rpp plus Fab30 (red) are indicated. Shift in the melt curve upon addition of V_2_Rpp or V_2_Rpp together with Fab30 (stabilizes an active conformation of β-arrestins) to β-arrestin-2 alone indicates formation of complexes, confirming our previous work^11^. **c**, quantitative analysis of various control conditions as well as binding of active β-arrestin-2 (plus V_2_Rpp and stabilized by Fab30) to G_αi_ (bound nonhydrolyzable GTP analog of GTP, GMP-PNP) as assessed using thermal structural stability assay. Derivative melting temperatures of the various reaction complexes were computed and plotted as indicated in the figure on y-axis. Each condition differed with regard to the presence of the components as indicated in the bar graphs. For all conditions, data were derived from three independent experiments. \**P*< 0.05, one way ANOVA with Bonferroni post hoc. Graphs show mean ± s.e.m.

**Extended Data Figure 3:**
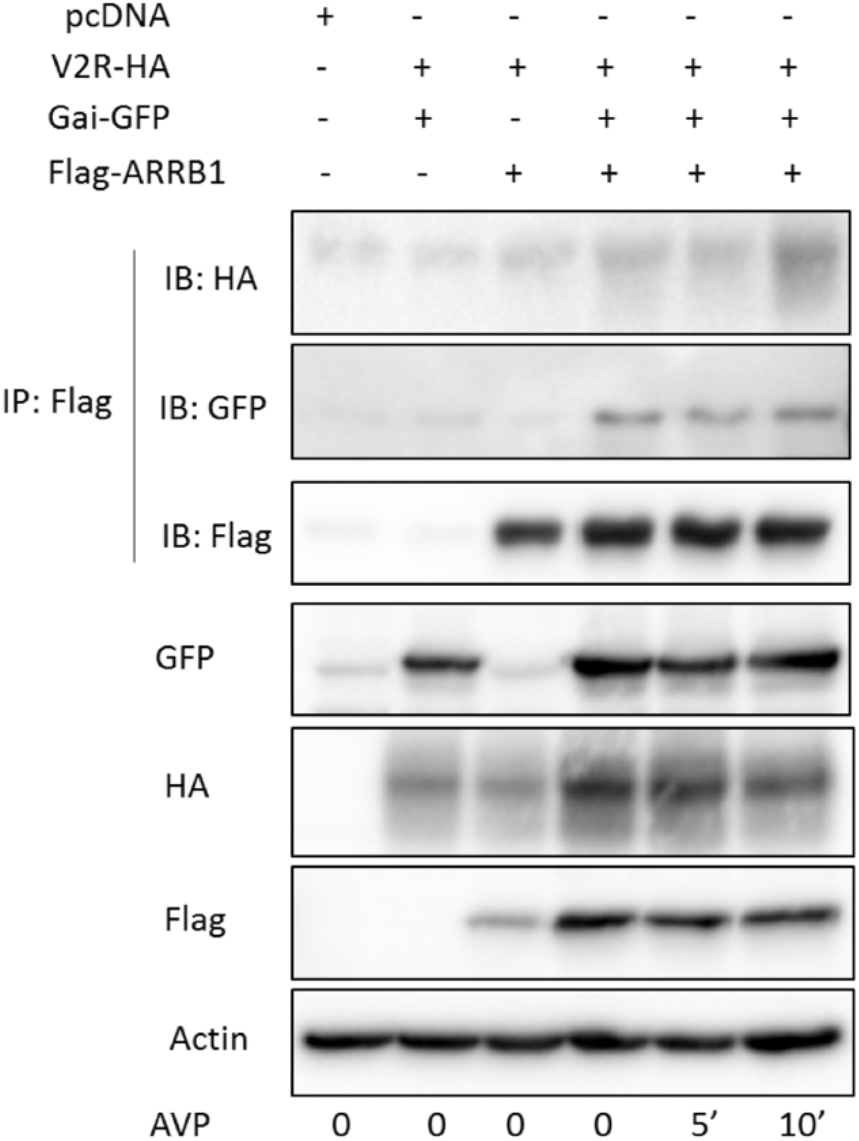
Immunoprecipitation of G_αi_:β-arrestin:V_2_R megaplex. HEK293 cells were transfected with the indicated plasmids and treated with AVP (500 nM) for the indicated duration. Co-transfection of G_αi_ and β-arrestin increased the expression of both proteins. Ligand treatment did not appreciably increase associated of G_αi_ and β-arrestin, which is explained by a high constitutive association required by the assay conditions and decreased granularity of signal relative to complex BRET (see extended Figure 6, where higher expression of assay components reduced agonist-induced signal). Data is representative of three separate experiments.

**Extended Data Figure 4:**
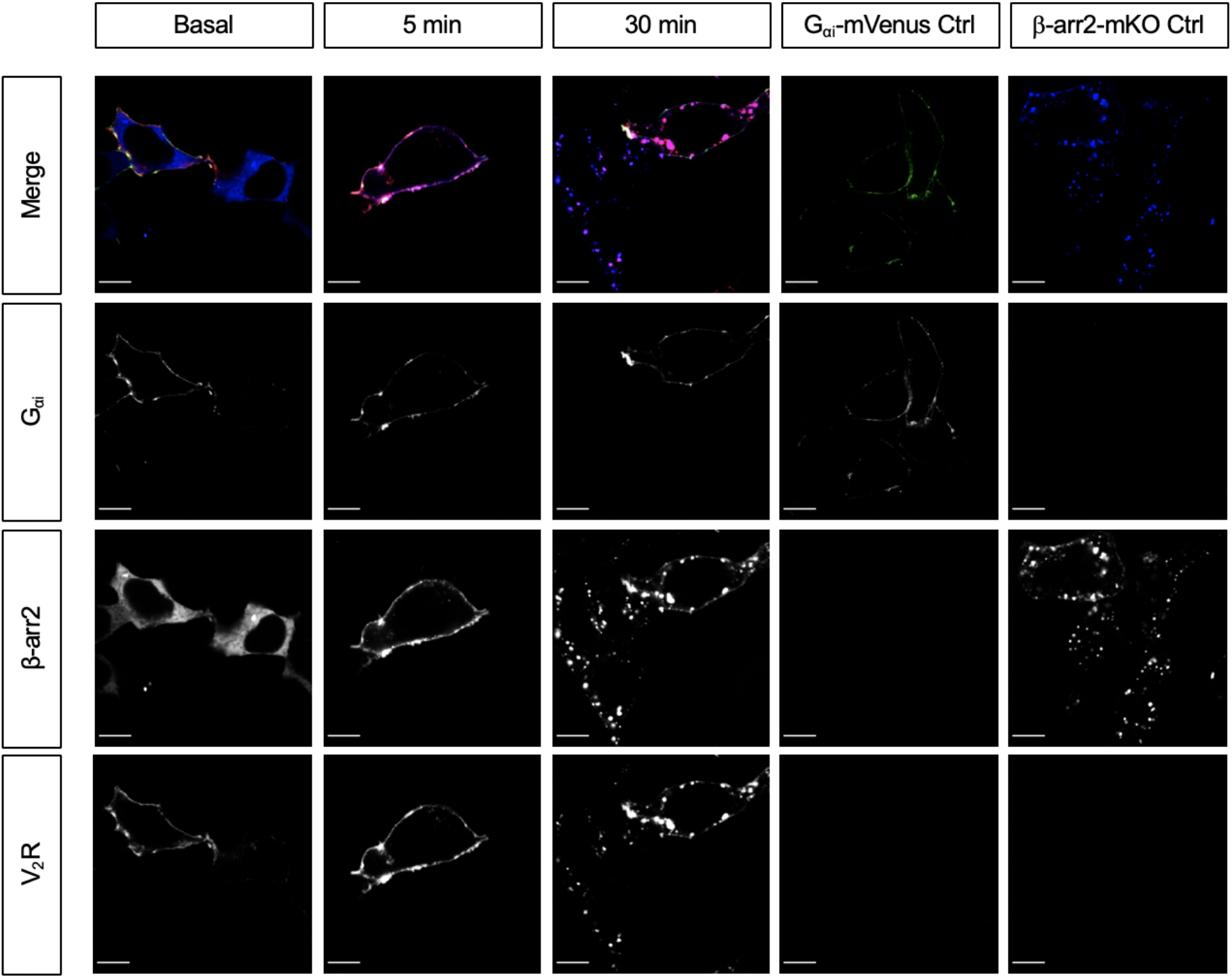
Single component controls to validate imaging parameters in Figure 2 of the main text. HEK293T cells transiently transfected with either all 3 components (G_αi_-mVenus, β-arrestin-2-mKO, Mars1-V_2_R) or single-colour controls and were then stimulated and fixed at various time points. Following this, the samples were imaged on a confocal microscope using identical parameters. All image adjustments were identical and consistent across all samples. Single-colour control samples (G_αi_-mVenus or β-arrestin-2-mKO alone) were used to verify that each imaging channel was only reporting on one component of the megaplex. Scale bars = 10 μm.

**Extended Data Figure 5:**
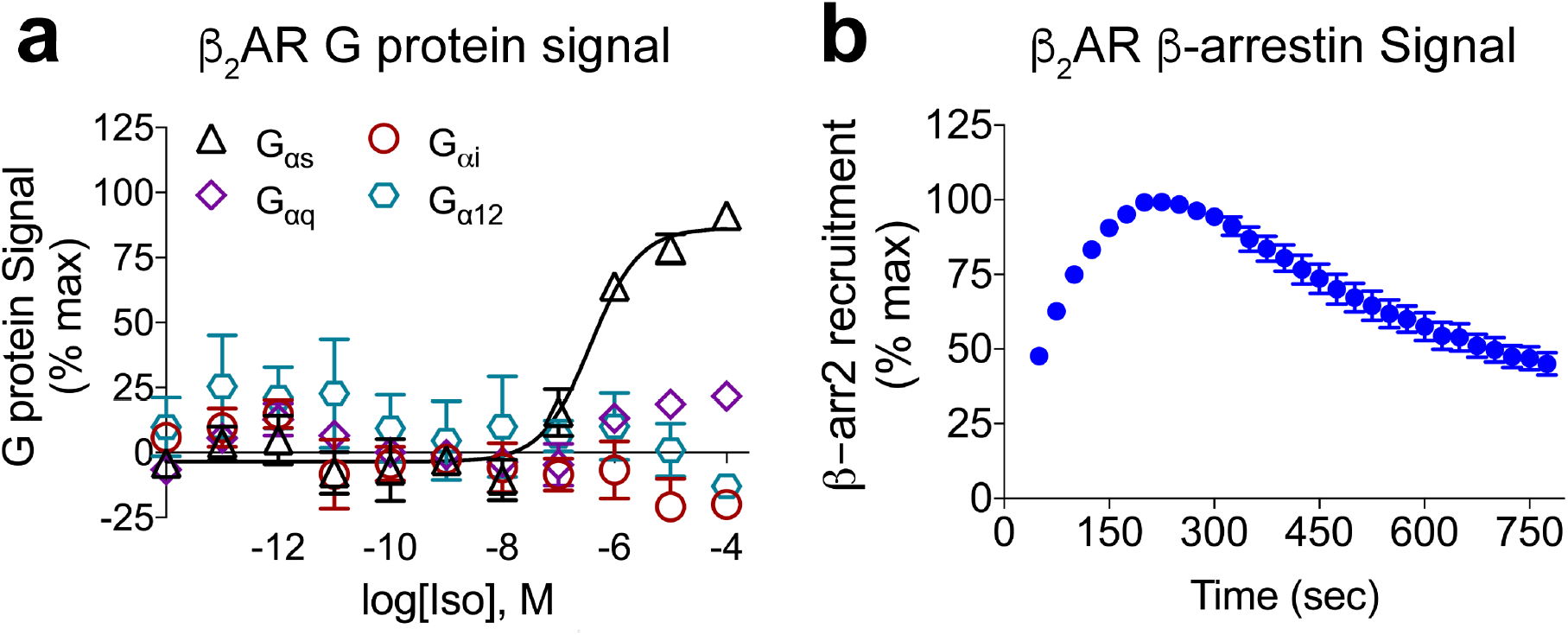
Assessment of β_2_AR-mediated G protein and β-arrestin signalling. **a**, Assessment of G protein signalling following agonist treatment of the β_2_AR in ‘ΔGsix’ HEK 293 cells transfected with the indicated G_α_ subunits and treated at the indicated concentration of isoproterenol. **b**, Assessment of β-arrestin-2 recruitment using luciferase complementation with β_2_AR-LgBiT and smBiT-β-arrestin-2 in WT HEK293T cells following treatment with isoproterenol (10 μM). n=3 per condition. Graphs show mean ± s.e.m.

**Extended Data Figure 6:**
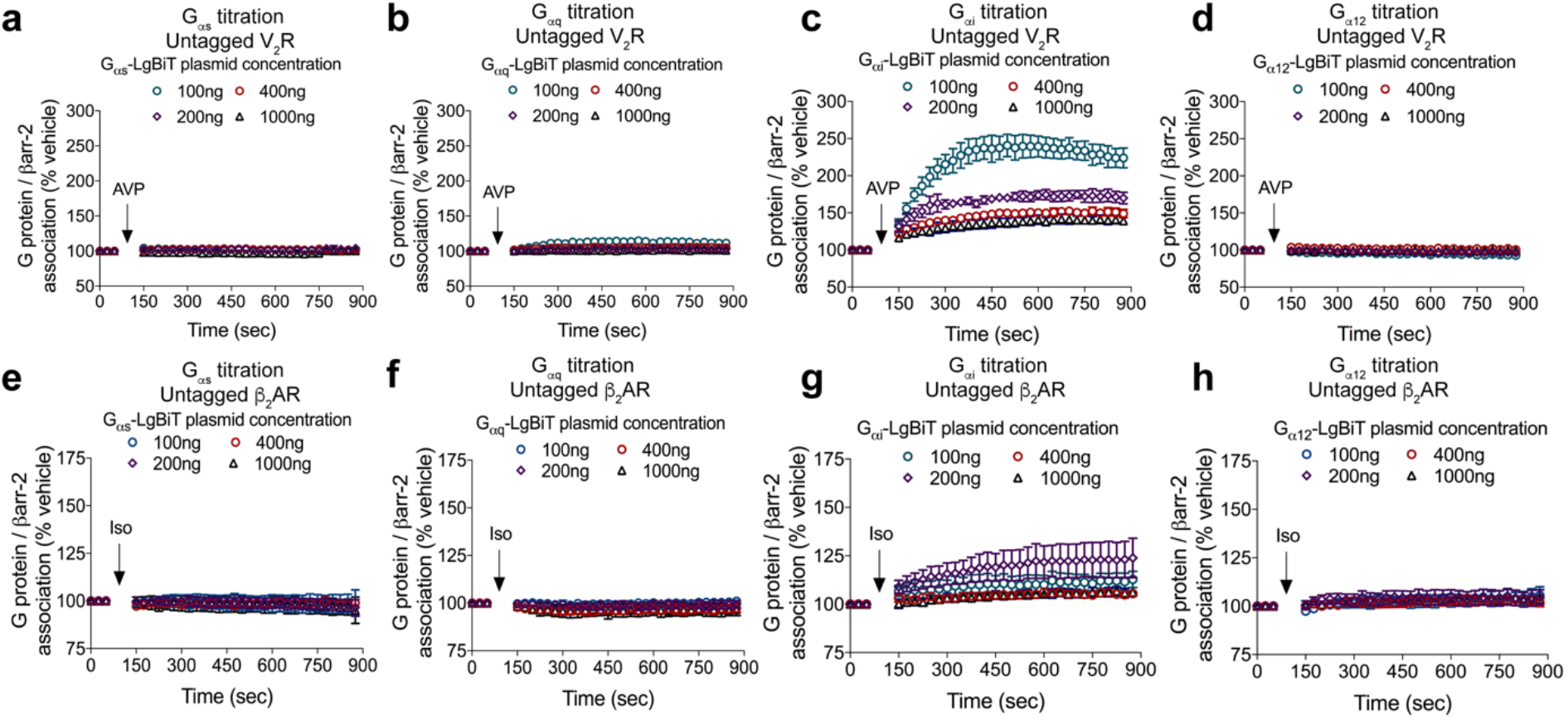
Increased overexpression of G_α_ decreases window for monitoring agonist-induced G_αs_, G_α12_, or G_αq_ association with β-arrestin-2. Split luciferase assay titration-response (cartoon shown in Fig. 3a) in HEK 293T cells quantifying agonist-stimulated G_α_-family proteins association with β-arrestin. HEK 293T cells transiently transfected with V_2_R and either 100 ng, 200ng, 400ng, or 1000ng of **a**, G_αs_-LgBiT, **b**, G_αq_-LgBiT, **c**, G_αi_-LgBiT, **d**, G_α12_-LgBiT expression vector and a constant 500 ng of SmBiT-β-arrestin-2 expression vector. Cells were treated with either vehicle or AVP (500 nM). Percentage signal above vehicle treatment is shown. Similarly, FLAG-β_2_AR and either 100, 200, 400, or 1000 ng of G_αs_-LgBiT **e**, G_αq_-LgBiT **f**, G_αi_-LgBiT **g**, G_α12_-LgBiT **h**, expression vector and 500 ng of SmBiT-β-arrestin-2 expression vector. Cells were treated with either vehicle or isoproterenol (10 uM). For panels **a,b**, n=2-3 per condition, for panel **c**, n=3-4 per condition, for panel **d**, n=3 per condition, for panel **e**, n=3-4 per condition, for panel **f**, n=2-3 per condition, for panel **g**, n=3-6 per condition, for panel **h**, n=3 per condition.

**Extended Data Figure 7:**
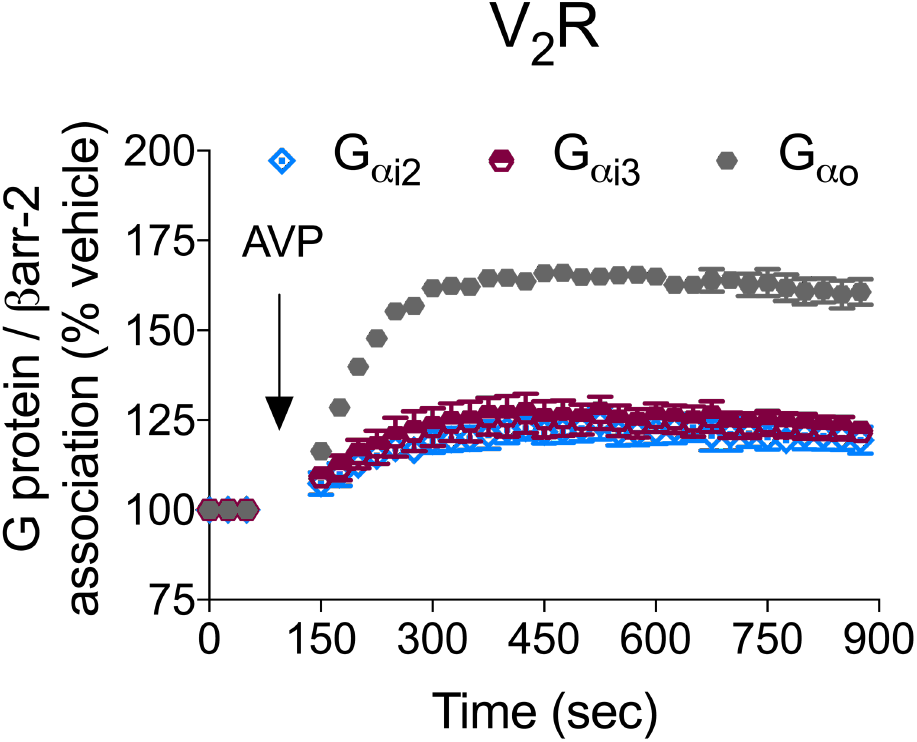
Other inhibitory G_α_ family members form a complex with β-arrestin following agonist treatment. HEK 293T cells were transfected with untagged V_2_R along with the indicated LgBiT-tagged G_αi_-family proteins (G_αi2_, G_αi3_, G_αo_), smBiT-β-arrestin-2, and treated with AVP (500 nM) or vehicle to quantify association with smBiT-β-arrestin-2. n=3 per condition. Graph shows mean ± s.e.m.

**Extended Data Figure 8:**
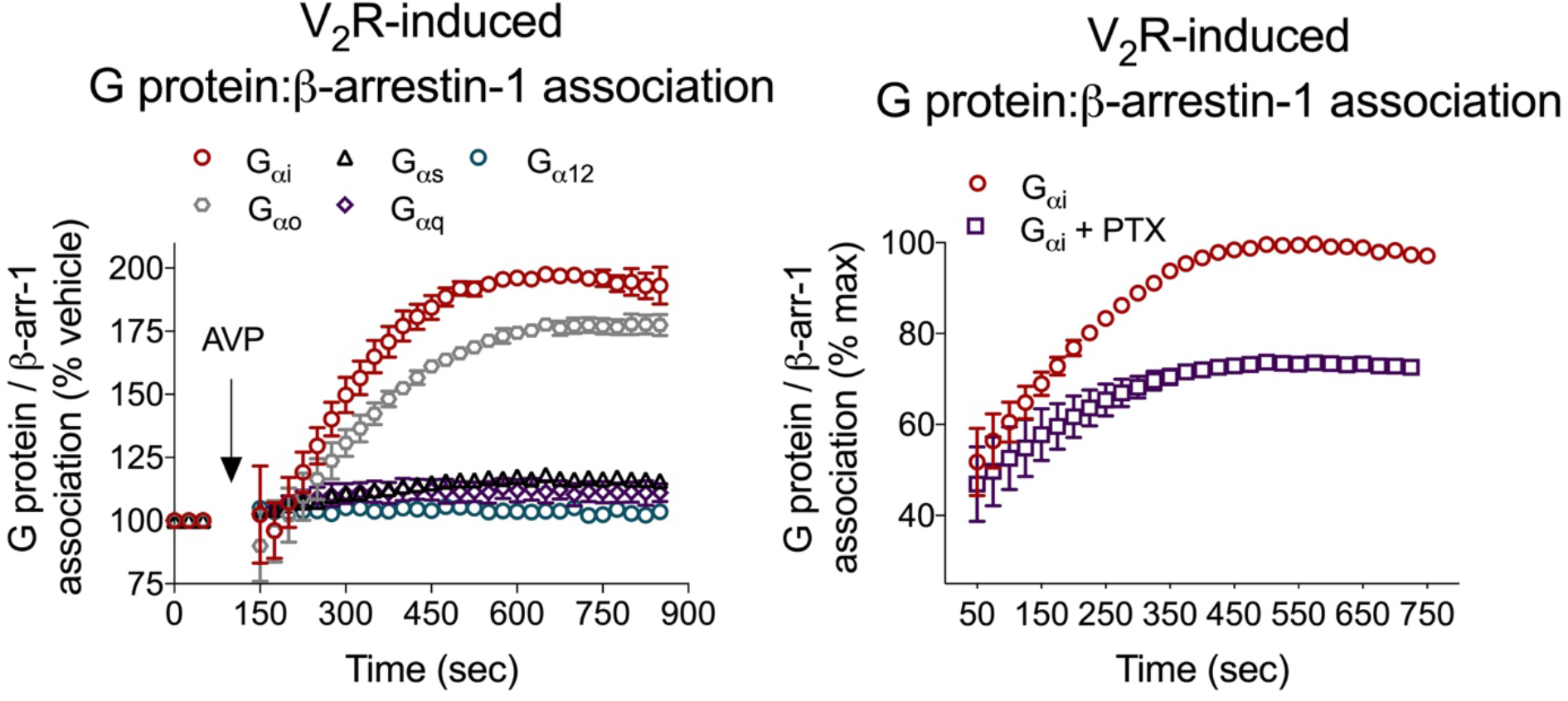
β-arrestin-1 shows a similar pattern to β-arrestin-2 of agonist-induced association with G_αi_-family that is *pertussis toxin* sensitive. **a**, HEK 293T cells were transiently transfected with V_2_R, smBiT-β-arrestin-1, and either LgBiT-tagged G_αi_, G_αo_, G_αq_, G_αs_, or G_α12_ and stimulated with AVP (500 nM). **b**, AVP-induced association of SmBiT-β-arrestin-1 and G_αi_-LgBiT was attenuated by *pertussis toxin* pretreatment (200 ng/mL). Luminescence values are normalized within well to signal prior to agonist treatment. n=3 replicates per condition. Graphs show mean ± s.e.m.

**Extended Data Figure 9:**
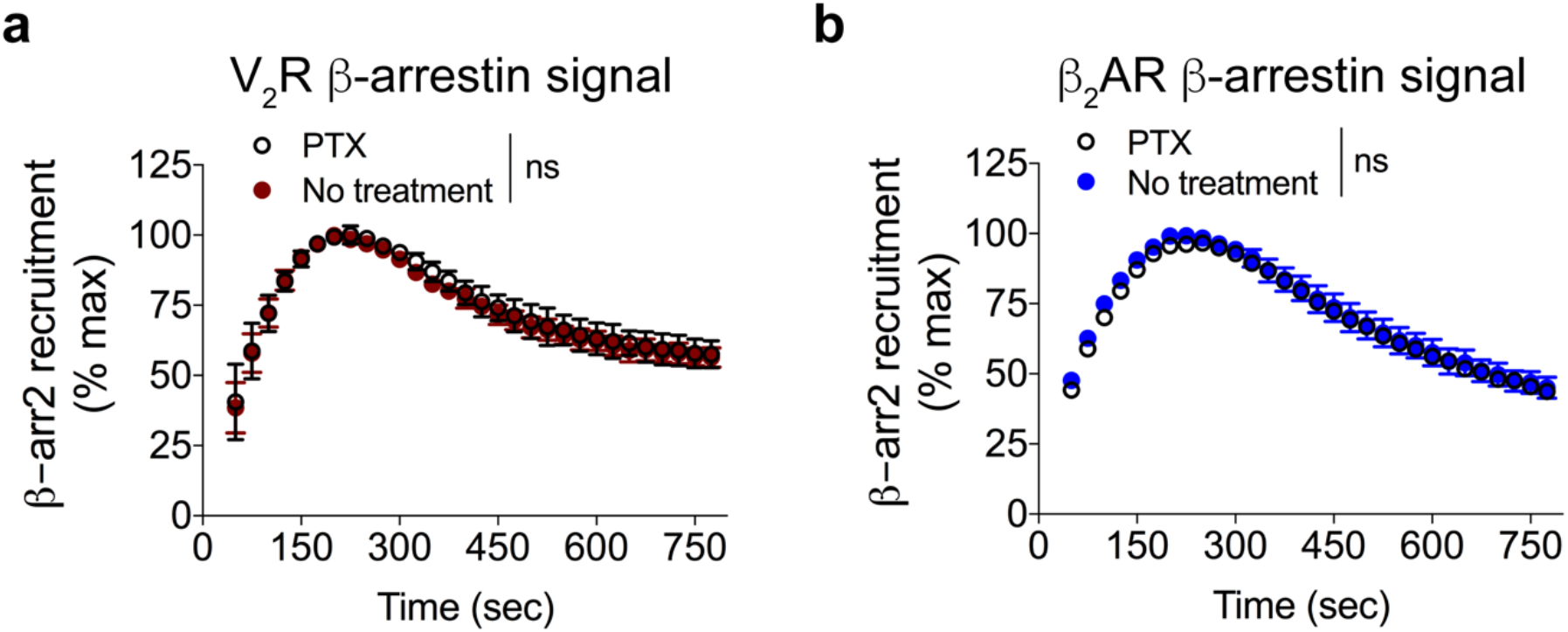
*Pertussis toxin* pretreatment does not affect agonist-induced β-arrestin-2 recruitment to either the V_2_R or the β_2_AR. HEK 293T cells were transiently transfected with smBiT-β-arrestin-2 and either V_2_R-LgBiT or β_2_AR-LgBiT. Cells were incubated with or without *pertussis toxin* (200 ng/mL). **a**, No effect of *pertussis toxin* pretreatment on AVP (500 nM)-induced β-arrestin-2 recruitment to V_2_R relative to non-treated controls. **b**, No effect of *pertussis toxin* pretreatment on isoproterenol (10 μM)-induced β-arrestin-2 recruitment to the β_2_AR relative to non-treated controls. n=3 per condition. Graphs show mean ± s.e.m.

**Extended Data Figure 10:**
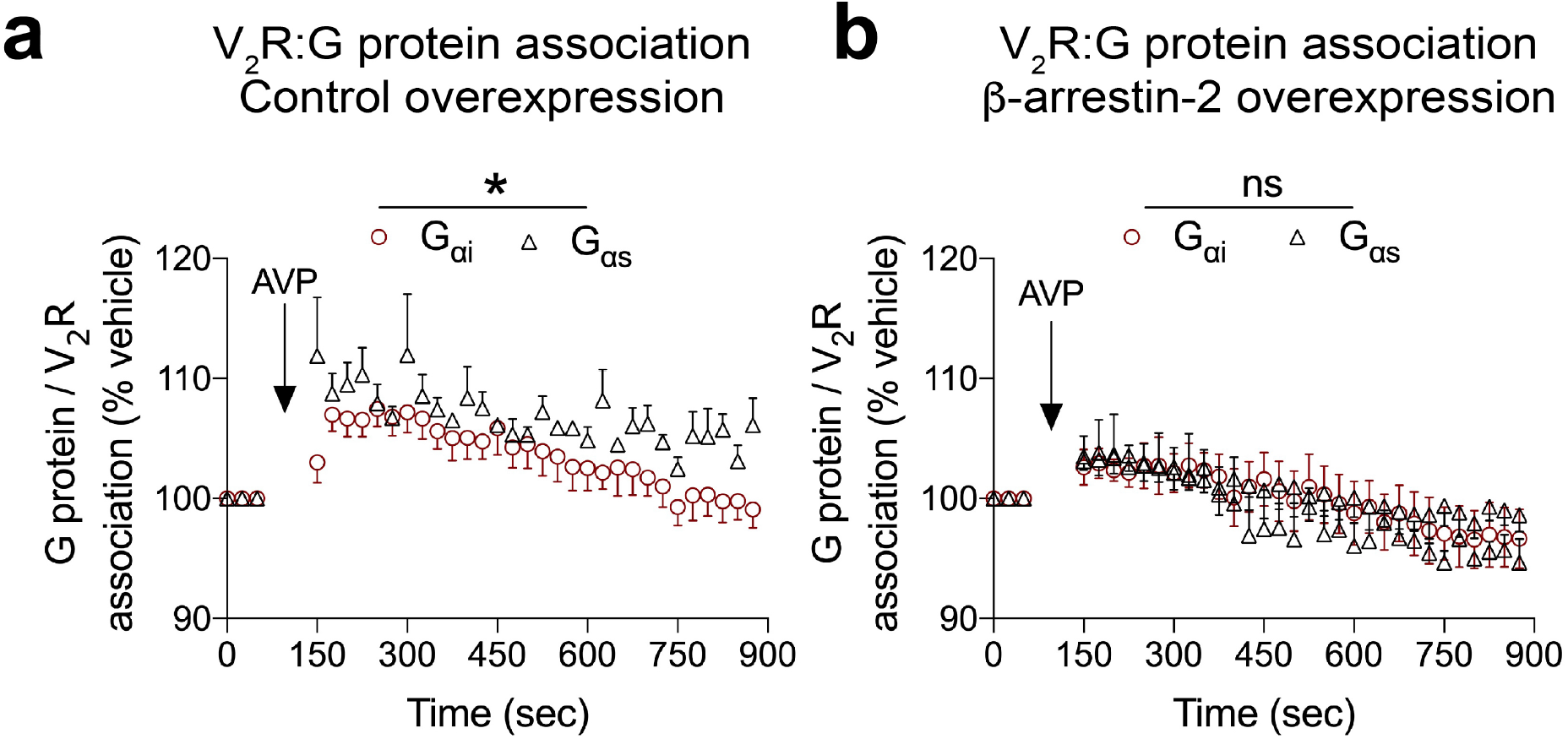
G_αi_ and G_αs_ are recruited to the V_2_R following agonist treatment. HEK 293T cells were transiently transfected with V_2_R-smBiT, either G_αi_-LgBiT or G_αs_-LgBiT, and either **a**, pcDNA or **b**, untagged β-arrestin-2. Cells were treated with AVP (500 nM), and association of V_2_R and the indicated G_α_ was measured by luminescence. Both G_αs_ and G_αi_ were recruited to V_2_R following AVP treatment, with the efficacy of the interaction significantly greater in the pcDNA control group, but not in the β-arrestin-2 overexpression group. \**P*<0.05, For panel **a**, n=3-5 per condition; for panel **s**, n=3-4 per condition. Graphs show mean ± s.e.m.

**Extended Data Figure 11:**
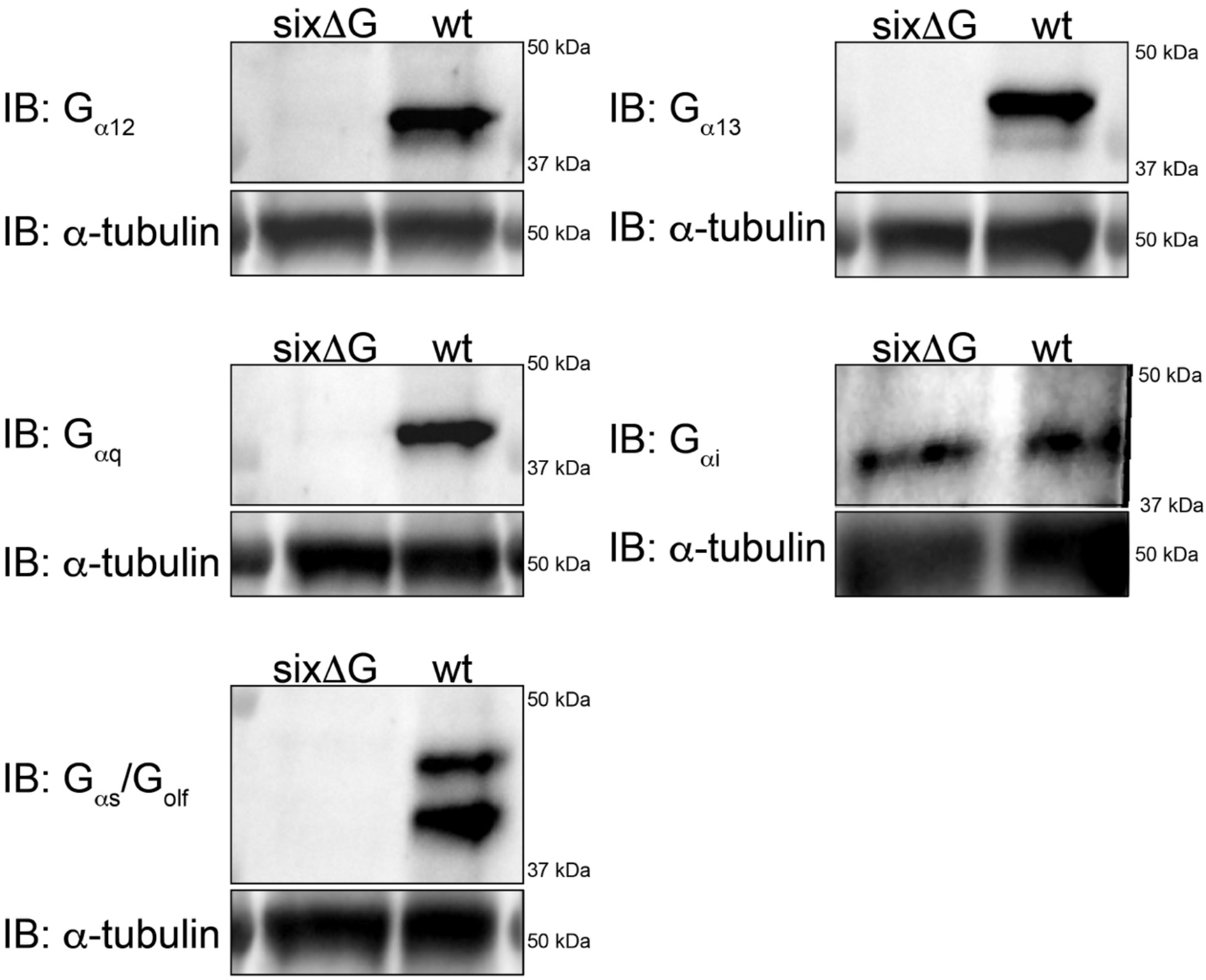
Validation of G_α_ protein knockout with CRISPR/Cas9. Immunoblot of lysates collected from previously described ‘ΔGsix’ HEK 293 cells shows only measurable G_αi_, but not other G_α_ protein family members. Lysates from parental WT HEK 293 cells are shown as a control.

## References

1 Gilman, A. G. G proteins: transducers of receptor-generated signals. Annu Rev Biochem 56, 615–649, doi:10.1146/annurev.bi.56.070187.003151 (1987).

2 Lohse, M. J., Benovic, J. L., Codina, J., Caron, M. G. & Lefkowitz, R. J. beta-Arrestin: a protein that regulates beta-adrenergic receptor function. Science 248, 1547–1550 (1990).

3 Thomsen, A. R. et al. GPCR-G Protein-beta-Arrestin Super-Complex Mediates Sustained G Protein Signaling. Cell 166, 907–919, doi:10.1016/j.cell.2016.07.004 (2016).

4 Wehbi, V. L. et al. Noncanonical GPCR signaling arising from a PTH receptor-arrestin-Gbetagamma complex. Proceedings of the National Academy of Sciences of the United States of America 110, 1530–1535, doi:10.1073/pnas.1205756110 (2013).

5 Oakley, R. H., Laporte, S. A., Holt, J. A., Barak, L. S. & Caron, M. G. Association of beta-arrestin with G protein-coupled receptors during clathrin-mediated endocytosis dictates the profile of receptor resensitization. The Journal of biological chemistry 274, 32248–32257 (1999).

6 Tohgo, A. et al. The stability of the G protein-coupled receptor-beta-arrestin interaction determines the mechanism and functional consequence of ERK activation. The Journal of biological chemistry 278, 6258–6267, doi:10.1074/jbc.M212231200 (2003).

7 Urizar, E. et al. CODA-RET reveals functional selectivity as a result of GPCR heteromerization. Nat Chem Biol 7, 624–630, doi:10.1038/nchembio.623 (2011).

8 Cotnoir-White, D. et al. Monitoring ligand-dependent assembly of receptor ternary complexes in live cells by BRETFect. Proceedings of the National Academy of Sciences of the United States of America 115, E2653–E2662, doi:10.1073/pnas.1716224115 (2018).

9 Dixon, A. S. et al. NanoLuc Complementation Reporter Optimized for Accurate Measurement of Protein Interactions in Cells. ACS Chem Biol 11, 400–408, doi:10.1021/acschembio.5b00753 (2016).

10 Renaud, J. P. et al. Biophysics in drug discovery: impact, challenges and opportunities. Nat Rev Drug Discov 15, 679–698, doi:10.1038/nrd.2016.123 (2016).

11 Shukla, A. K. et al. Structure of active beta-arrestin-1 bound to a G-protein-coupled receptor phosphopeptide. Nature 497, 137–141, doi:10.1038/nature12120 (2013).

12 Irannejad, R. et al. Functional selectivity of GPCR-directed drug action through location bias. Nat Chem Biol 13, 799–806, doi:10.1038/nchembio.2389 (2017).

13 Eichel, K. et al. Catalytic activation of beta-arrestin by GPCRs. Nature 557, 381–386, doi:10.1038/s41586-018-0079-1 (2018).

14 West, R. E., Jr., Moss, J., Vaughan, M., Liu, T. & Liu, T. Y. Pertussis toxin-catalyzed ADP-ribosylation of transducin. Cysteine 347 is the ADP-ribose acceptor site. The Journal of biological chemistry 260, 14428–14430 (1985).

15 Pack, T. F., Orlen, M. I., Ray, C., Peterson, S. M. & Caron, M. G. The dopamine D2 receptor can directly recruit and activate GRK2 without G protein activation. J Biol Chem 293, 6161–6171, doi:10.1074/jbc.RA117.001300 (2018).

16 Luttrell, L. M. et al. Manifold roles of beta-arrestins in GPCR signaling elucidated with siRNA and CRISPR/Cas9. Sci Signal 11, doi:10.1126/scisignal.aat7650 (2018).

17 Wan, Q. et al. Mini G protein probes for active G protein-coupled receptors (GPCRs) in live cells. The Journal of biological chemistry 293, 7466–7473, doi:10.1074/jbc.RA118.001975 (2018).

18 Pearson, G. et al. Mitogen-activated protein (MAP) kinase pathways: regulation and physiological functions. Endocr Rev 22, 153–183, doi:10.1210/edrv.22.2.0428 (2001).

19 Smith, J. S., Lefkowitz, R. J. & Rajagopal, S. Biased signalling: from simple switches to allosteric microprocessors. Nat Rev Drug Discov 17, 243–260, doi:10.1038/nrd.2017.229 (2018).

20 Grundmann, M. et al. Lack of beta-arrestin signaling in the absence of active G proteins. Nat Commun 9, 341, doi:10.1038/s41467-017-02661-3 (2018).

21 O’Hayre, M. et al. Genetic evidence that beta-arrestins are dispensable for the initiation of beta2-adrenergic receptor signaling to ERK. Sci Signal 10, doi:10.1126/scisignal.aal3395 (2017).

22 Wang, J. et al. Galphai is required for carvedilol-induced beta1 adrenergic receptor beta-arrestin biased signaling. Nat Commun 8, 1706, doi:10.1038/s41467-017-01855-z (2017).

23 Hawes, B. E., van Biesen, T., Koch, W. J., Luttrell, L. M. & Lefkowitz, R. J. Distinct pathways of Gi-and Gq-mediated mitogen-activated protein kinase activation. The Journal of biological chemistry 270, 17148–17153 (1995).

24 Hordijk, P. L., Verlaan, I., van Corven, E. J. & Moolenaar, W. H. Protein tyrosine phosphorylation induced by lysophosphatidic acid in Rat-1 fibroblasts. Evidence that phosphorylation of map kinase is mediated by the Gi-p21ras pathway. The Journal of biological chemistry 269, 645–651 (1994).

25 Smith, J. S. & Rajagopal, S. The beta-Arrestins: Multifunctional Regulators of G Protein-coupled Receptors. The Journal of biological chemistry 291, 8969–8977, doi:10.1074/jbc.R115.713313 (2016).

26 Violin, J. D. et al. Selectively engaging beta-arrestins at the angiotensin II type 1 receptor reduces blood pressure and increases cardiac performance. The Journal of pharmacology and experimental therapeutics 335, 572–579, doi:10.1124/jpet.110.173005 (2010).

27 Strachan, R. T. et al. Divergent transducer-specific molecular efficacies generate biased agonism at a G protein-coupled receptor (GPCR). The Journal of biological chemistry 289, 14211–14224, doi:10.1074/jbc.M114.548131 (2014).

28 Smith, J. S. et al. Biased agonists of the chemokine receptor CXCR3 differentially control chemotaxis and inflammation. Sci Signal 11, doi:10.1126/scisignal.aaq1075 (2018).

29 Inoue, A. et al. TGFalpha shedding assay: an accurate and versatile method for detecting GPCR activation. Nat Methods 9, 1021–1029, doi:10.1038/nmeth.2172 (2012).

30 Gregorio, G. G. et al. Single-molecule analysis of ligand efficacy in beta2AR-G-protein activation. Nature 547, 68–73, doi:10.1038/nature22354 (2017).

31 Ma, Z., Chalkley, R. J. & Vosseller, K. Hyper-O-GlcNAcylation activates nuclear factor kappa-light-chain-enhancer of activated B cells (NF-kappaB) signaling through interplay with phosphorylation and acetylation. The Journal of biological chemistry 292, 9150–9163, doi:10.1074/jbc.M116.766568 (2017).

32 Snyder, J. C. et al. A rapid and affordable screening platform for membrane protein trafficking. BMC Biol 13, 107, doi:10.1186/s12915-015-0216-3 (2015).

